# Statistically unbiased prediction enables accurate denoising of voltage imaging data

**DOI:** 10.1101/2022.11.17.516709

**Authors:** Minho Eom, Seungjae Han, Gyuri Kim, Eun-Seo Cho, Jueun Sim, Pojeong Park, Kang-Han Lee, Seonghoon Kim, Márton Rózsa, Karel Svoboda, Myunghwan Choi, Cheol-Hee Kim, Adam E. Cohen, Jae-Byum Chang, Young-Gyu Yoon

**Affiliations:** School of Electrical Engineering, Korea Advanced Institute of Science and Technology, Daejeon, Republic of Korea; Department of Materials Science and Engineering, Korea Advanced Institute of Science and Technology, Daejeon, Korea; Department of Chemistry and Chemical Biology, Harvard University, Cambridge, United States; Department of Biology, Chungnam National University, Daejeon, South Korea; School of Biological Sciences, Seoul National University, Seoul, Republic of Korea; Institute of Molecular Biology and Genetics, Seoul National University, Seoul, Republic of Korea; Allen Institute for Neural Dynamics, Seattle, WA, USA; Department of Physics, Harvard University, Cambridge, United States; KAIST Institute for Health Science and Technology, Daejeon, Republic of Korea

**Author notes:** These authors contributed equally: Minho Eom and Seungjae Han. Corresponding Author: Young-Gyu Yoon.

## Abstract

Here we report SUPPORT (Statistically Unbiased Prediction utilizing sPatiOtempoRal information in imaging daTa), a self-supervised learning method for removing Poisson-Gaussian noise in voltage imaging data. SUPPORT is based on the insight that a pixel value in voltage imaging data is highly dependent on its spatially neighboring pixels in the same time frame, even when its temporally adjacent frames do not provide useful information for statistical prediction. Such spatiotemporal dependency is captured and utilized to accurately denoise voltage imaging data in which the existence of the action potential in a time frame cannot be inferred by the information in other frames. Through simulation and experiments, we show that SUPPORT enables precise denoising of voltage imaging data while preserving the underlying dynamics in the scene.

## MAIN

Recent advancements in voltage imaging and calcium imaging have enabled the recording of the population activity of neurons at an unprecedented throughput, which opens up the possibility of a system-level understanding of neuronal circuits^1–3^. To investigate the causality within neuronal activities, it is essential to record the activities of the population of neurons with high temporal precision. Unfortunately, the inherent limitation in the maximum number of photons that can be collected from a sample in a given time interval dictates the inherent trade-offs between imaging speed and signal-to-noise ratio (SNR)^4,5^. In other words, increasing the temporal resolution in functional imaging data inevitably results in a decrease in the SNR. The decrease in SNR not only hinders the accurate detection of the neurons’ location but also compromises the timing precision of the detected temporal events, which nullifies the increase in temporal resolution. Fortunately, all functional imaging data have high inherent redundancy in the sense that each frame in a dataset is nearly identical to other frames apart from noise, which offers an opportunity to denoise or distinguish the signal from the noise in the data^6–8^.

Denoising is a type of signal processing that attempts to extract underlying signals from noisy observations based on prior knowledge of the signal and the noise^9^. Importantly, the fundamental property of noise—randomness—does not allow for exact recovery of the signal, so we can only reduce statistical variance at the cost of increasing statistical bias (i.e., an absolute deviation between the mean denoising outcome and the ground truth). In other words, denoising is a statistical estimation of the most probable value based on our prior statistical knowledge of the signal and the noise. Unfortunately, for any given noisy observation, the exact corresponding probability distribution functions (PDFs) of the signal and the noise are almost never known. Therefore, all denoising algorithms start by setting the signal model (i.e., PDF of signal) and noise model (i.e., PDF of noise), either explicitly or implicitly, and their accuracy determines the denoising performance to a great extent.

The most common approach starts with applying linear transforms, such as the Fourier transform and the wavelet transform, to noisy observations^10,11^. Then, a certain set of coefficients that corresponds to a small vector space is preserved, while others are attenuated to reduce statistical variance. This is based on a signal model in which the signal is a random variable drawn from the small vector space, whereas noise is drawn from the entire vector space. An implicit yet important assumption here is that the basis used for the linear transform maps the signal component sharply onto a relatively small and known set of coefficients. When the assumption is not met, denoising leads to a distortion of signals or an increase in statistical bias. Such bias can be reduced by loosening the assumption (e.g., the signal is drawn from a larger vector space), but then the variance is increased.

Therefore, building a good signal model that is strong enough to reject noise while being accurate enough to avoid bias is the most critical step in denoising. Previous efforts have focused on finding a handcrafted basis that empirically matches the given data^12^. Some have shown higher general applicability than others^13^, but no universal basis that performs well across different types of data has been found, mainly because of the differences in their signal models and noise models^14^. This has led to the idea of using a basis that is learned directly from the dataset for denoising^6,15,16^. However, these methods still suffer from high bias, as their ability to reduce variance relies on the strong assumption that the data can be represented as a linear summation of a small number of learned vectors.

Recently, the convolutional network has emerged as a strong alternative to existing learning-based image denoising algorithms^17^. The high representational power of convolutional networks allows for learning nearly arbitrary signal models in the image domain, which result in low bias in denoising outcomes without sacrificing variance^18^. Owing to its high representational power and the high inherent redundancy in functional imaging data, convolutional networks have shown enormous success in denoising functional imaging data^7,8^. As a key aspect, these methods learn the signal model from noisy data in a self-supervised manner^19–22^, so the need for “clean” images is alleviated.

Both DeepCAD-RT^7^ and DeepInterpolation^8^ are based on the assumption that the underlying signal in any two consecutive frames in a video can be considered the same, whereas the noise is independent when the imaging speed is sufficiently higher than the dynamics of the fluorescent reporter^7,8^; the networks are trained to predict the “current” frame using the past and future frames as the input. Unfortunately, this assumption breaks down when the imaging speed is not sufficiently faster than the dynamics, and the bias in the denoising outcome is increased. This is becoming increasingly prevalent due to the development of voltage indicators^23–28^ and calcium indicators with extremely fast dynamics^29^. In that regard, the question that naturally follows is how we can implement an accurate statistical model that allows us to accurately predict each pixel value under such conditions.

To this end, we propose SUPPORT (Statistically Unbiased Prediction utilizing sPatiOtempoRal information in imaging daTa), a self-supervised denoising method for functional imaging data that is robust to fast dynamics in the scene compared to the imaging speed. SUPPORT is based on the insight that a pixel value in functional imaging data is highly dependent on its spatially neighboring pixels in the same time frame, even when its temporally adjacent frames fail to provide useful information for statistical prediction. By learning and utilizing the spatiotemporal dependence among the pixels, SUPPORT can accurately denoise voltage imaging data in which the existence of the action potential in a time frame cannot be inferred by the information in other frames. We demonstrate the capability of SUPPORT using diverse voltage imaging datasets acquired using Voltron1, Voltron2, paQuasAr3-s, QuasAr6a, zArchon1, and BeRST1. The analysis of the voltage imaging data with simultaneous electrophysiology recording shows that our method preserves the shape of the spike while reducing the statistical variance in the signal. We also show that SUPPORT can be used for denoising time-lapse fluorescence microscopy images of *Caenorhabditis elegans* (*C. elegans*), in which the imaging speed is not faster than the locomotion of the worm, as well as static volumetric images of *Penicillium* and mouse embryos. SUPPORT is exceptionally compelling for denoising voltage imaging and time-lapse imaging data, and is even effective for denoising calcium imaging data. Finally, we developed software with a graphical user interface (GUI) for running SUPPORT to make it available to the wider community.

## RESULTS

### General working principle of SUPPORT

The main philosophy of SUPPORT is to perform denoising based on an unbiased statistical prediction model by exploiting all available information in both spatial and temporal domains (Fig. 1a). A functional imaging dataset ***y*** is considered a realization of a random variable that is drawn from *p*(***y***) = *p*(***x***)*p*(***n***|***x***), where ***x*** and ***n*** are the clean signal and the zero-mean Poisson-Gaussian additive noise, respectively (i.e., ***y*** = ***x*** + ***n***). In this setting, the noise in each pixel is independent in both time and space (i.e., ∀(*i, k*) ≠ (*j, l*), *p*(*n*_i,k_) = *p*(*n*_i,k_|*n*_j,l_), where *i, j* and *k, l* are temporal and spatial indices, respectively, where the signal is not (i.e., ∀(*i, k, j, l*), *p*(*x*_i,k_) ≠ *p*(*x*_i,k_|*x*_j,l_)). The dependency among *x*_i,k_ encodes the spatiotemporal structure of the data ***x*** (i.e., *p*(***x***)), which can be learned using a statistical prediction model, whereas the spatiotemporal independence of ***n*** makes it impossible to predict. The prediction model can be implemented as a neural network that predicts a pixel value *x*_i,k_ using its spatiotemporal neighboring pixel values by solving the following optimization problem^19,20^:

**Fig. 1:**
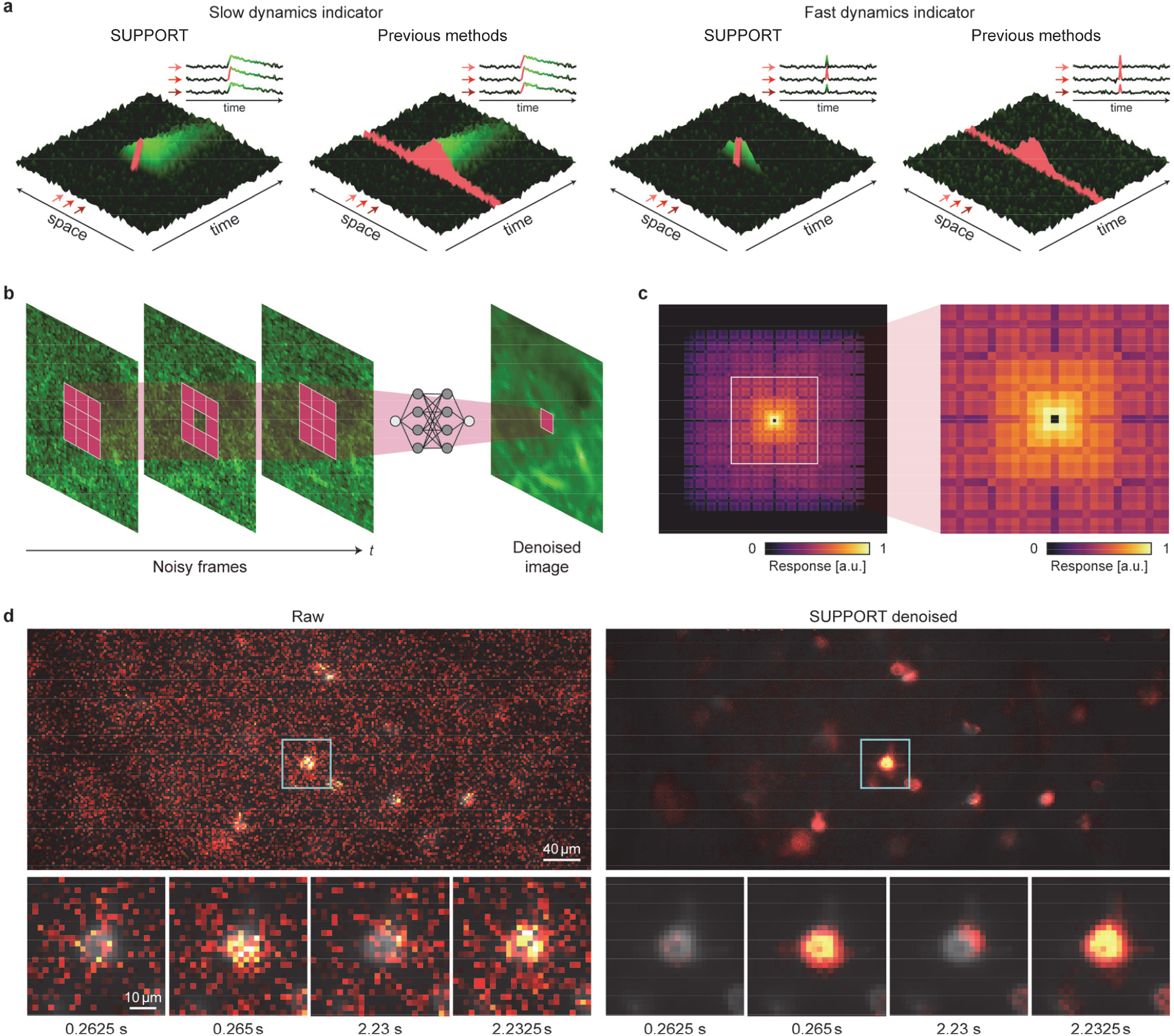
SUPPORT can be applied to functional imaging data with a fast dynamics indicator. **a**, SUPPORT’s self-supervised learning scheme and previous methods that exploit temporally adjacent frames for denoising functional imaging data with slow and fast dynamics indicators. Functional imaging data are represented by green and red surfaces, which indicate the receptive field and prediction target area, respectively. **b**, Noisy frames are fed into the SUPPORT network and output the denoised image. Red tiles indicate the receptive field of the SUPPORT network, which utilizes spatially adjacent pixels in the same frame. **c**, Impulse response of the SUPPORT network on the current frame. Magnified view is presented on the right side. Response value of the center pixel is 0, which forces the network to predict the center pixel without utilizing it. **d**, In vivo population voltage imaging data. Left: Raw data. Right: SUPPORT denoised data. Baseline and activity components are decomposed from raw data and SUPPORT denoised data. The baseline component with gray colormap and activity component with hot colormap are overlaid. Magnified views of the boxed regions are presented below at the time points near spikes. Consecutive frames of two spikes (t = 0.265 s, 2.2325 s).

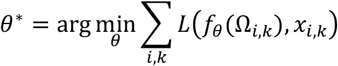

where *L*(·,·) is the loss function defined as the *L*^p^-distance between the inputs, *f*_0_ denotes the neural network parameterized by *θ*, and Ω_i,k_ denotes the spatiotemporal neighboring pixels of *y*_i,k_ excluding itself. Evaluating this loss function requires the ground truth ***x***, which is inaccessible, but the zero-mean property of the noise allows us to replace *x*_i,k_ with *y*_i,k_ for self-supervised training^20^:

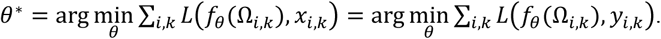

For the implementation of the network *f*_0_(Ω_i,k_), we devised a network architecture that automatically satisfies the requirements (Fig. 1b, c, Supplementary Fig. 1, 2): for the prediction *x*_i,k_, its spatiotemporal neighbor Ω_i,k_ excluding *y*_i,k_ is taken as the input while preserving the spatial invariance. The current frame *y*_i_ is fed into a convolutional network that has a zero at the center of the impulse response (Fig. 1b,c); the zero at the center of the impulse response indicates that the pixel value *y*_i,k_ cannot affect the network’s prediction of *x*_i,k_^19,30^, which is attained by convolution layers and dilated convolution layers with zeros at the center of the kernels. These layers offer a fractal-shaped receptive field that grows exponentially with depth, enabling the network to integrate information from a large number of neighboring pixels (Supplementary Fig. 2). In addition, temporally adjacent frames are fed into a U-Net^31^ to extract the available information from the temporally adjacent frames (Supplementary Fig. 1). The outputs from the two convolutional networks are integrated by the following convolutional layers. This architecture “forces” the network to make a prediction 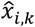 by utilizing its spatiotemporal neighbor Ω_i,k_ excluding *y*_i,k_ (i.e., 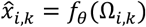).

The major difference between SUPPORT and DeepCAD-RT^7^ or DeepInterpolation^8^, which can also denoise functional imaging data through self-supervised learning, is that DeepCAD-RT and DeepInterpolation learn to predict a frame given temporally adjacent other frames, whereas SUPPORT learns to predict each pixel value by exploiting the information available from both temporally adjacent frames and spatially adjacent pixels in the same time frame. When the imaging speed is not sufficiently faster than the dynamics in the scene (Fig. 1a), the signal at different time points becomes nearly independent (e.g., the existence of the action potential in a time frame cannot be inferred from the information in other frames). In such a case, the major assumptions of the signal models in DeepCAD-RT and DeepInterpolation are violated, which leads to high bias in the denoising outcome. In comparison, SUPPORT relies on the spatiotemporal pixel-level dependence of the signal rather than frame-level dependence, and each pixel value is estimated based on all available information, including its spatially adjacent pixels in the same time frame. This prevents the violation of the assumption in the signal model, even when the imaging speed is insufficient; hence, a low bias result can be obtained without sacrificing variance.

### Performance validation on simulated data

For the quantitative evaluation of SUPPORT’s performance, we first validated it on synthetic voltage imaging data, which were generated using a NAOMi simulator^32^. We generated multiple datasets with a frame rate of 500 Hz with different spike widths, ranging from 1 ms to 9 ms^33^, to verify how the performance of SUPPORT changes as the dependence among the activity in adjacent frames is diminished. The simulation parameters, including spike frequency, dF/F_0_, noise level, and level of subthreshold activity, were chosen to match the experimental voltage imaging data acquired using Voltron^23^ (Methods). Finally, Poisson and Gaussian noise were added to the generated videos. Further details can be found in the Methods section.

We applied SUPPORT, DeepCAD-RT^7^, and penalized matrix decomposition (PMD)^6^ to the synthetic datasets and compared the results. The signals were separated from the backgrounds in the denoised videos (Methods) to compare their accuracy in recovering the time-varying signal (Fig. 2a, Supplementary Video 1). Qualitative comparisons of the results from the dataset with a spike width of 3 ms showed that the denoising outcome from SUPPORT was nearly identical to the ground truth. DeepCAD-RT successfully reduced the variance in the video, but also attenuated the neuronal activity. This was expected because the method was designed for removing noise in calcium imaging data, which has much slower dynamics. PMD showed better performance in preserving neuronal activities, in part because it did not discard the current frame for denoising, but it introduced visible artifacts in the images.

**Fig. 2:**
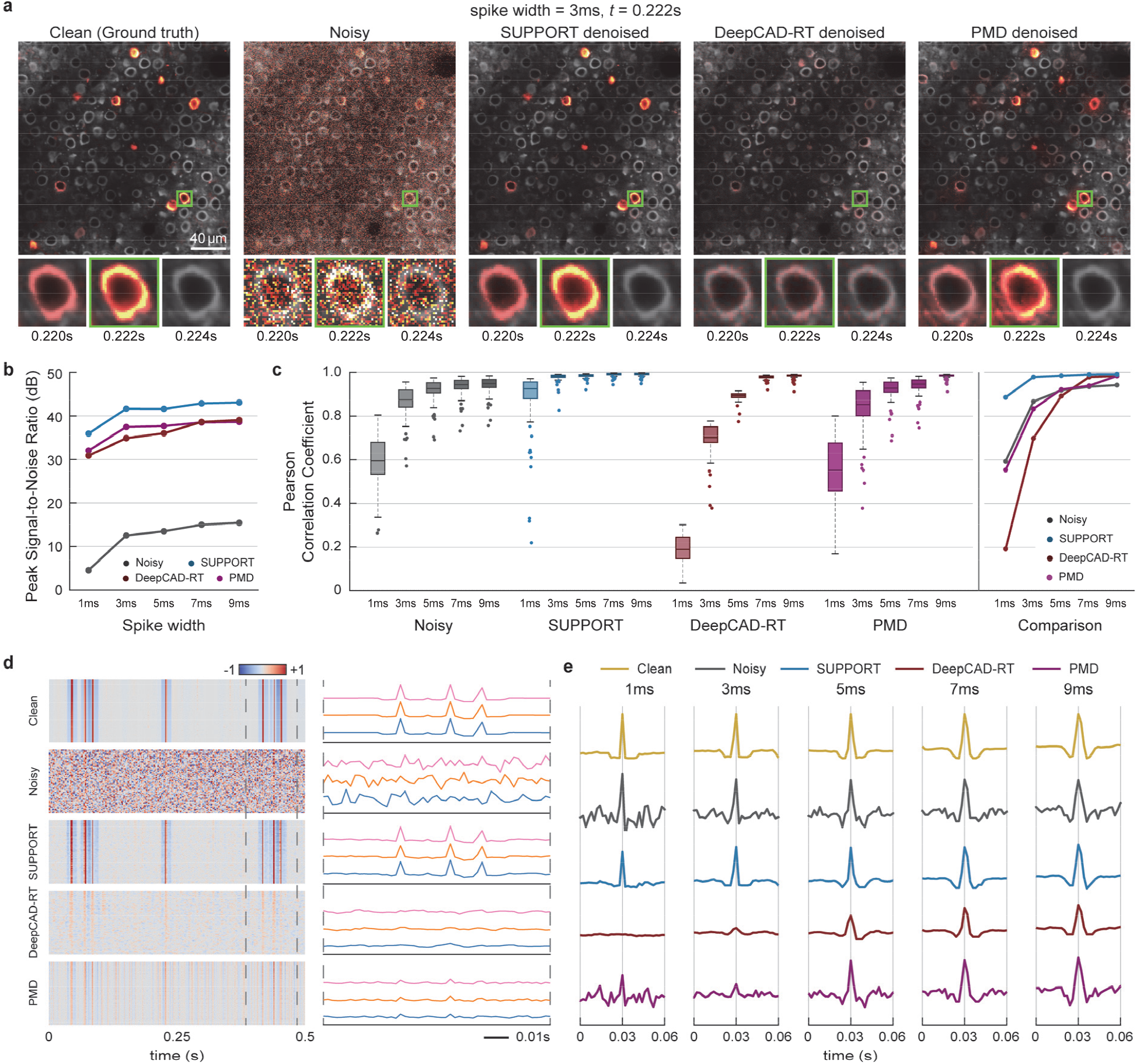
Performance validation on simulated data. **a**, Synthetic population voltage imaging data. From left to right: Clean, noisy, SUPPORT, DeepCAD-RT, and PMD denoised data. Baseline and activity components are decomposed from the data. The baseline component with a gray colormap and activity component with a hot colormap are overlaid. Magnified views of the boxed regions are presented underneath with the consecutive frames of the spiking event (t = 0.222 s). **b**, PSNR of the baseline-corrected data before and after denoising data with different spike widths. Clean data were used as the ground truth for PSNR calculation. **c**, Left: Box-and-whisker plot showing Pearson correlation coefficients before and after denoising data with different spike widths. Right: Line chart showing average Pearson correlation coefficients before and after denoising data with different spike widths; *N* = 116. **d**, Single pixel fluorescence traces extracted from baseline-corrected data. From top to bottom: Clean, noisy, SUPPORT, DeepCAD-RT, and PMD denoised data. Left: Each single pixel trace occupies each row. Right: Three representative single pixel traces visualized with different colors. **e**, Single cell fluorescence traces near spiking event extracted from baseline-corrected data. From top to bottom: Clean, noisy, SUPPORT, DeepCAD-RT, and PMD denoised data. From left to right: Changing spike widths of 1 ms, 3 ms, 5 ms, 7 ms, and 9 ms.

To quantify the performance of each denoising method, we calculated the peak signal-to-noise ratio (PSNR) of the denoised videos and calculated the Pearson correlation coefficient between the voltage traces extracted from the clean video and the denoised video. The voltage traces were extracted from 116 cells (Methods). In terms of PSNR, all methods showed significant enhancements compared to noisy images for every spike width (Fig. 2b): Noisy (1 ms: 4.57 dB, 9 ms: 15.43 dB), SUPPORT (1 ms: 35.94 dB, 9 ms: 43.08 dB), DeepCAD-RT (1 ms: 30.90 dB, 9 ms: 39.05 dB), and PMD (1 ms: 32.07 dB, 9 ms: 38.61 dB). However, in terms of the Pearson correlation coefficient, only SUPPORT (1 ms: 0.885, 9 ms: 0.991) showed enhancement compared to noisy images (1 ms: 0.593, 9 ms: 0.942) for every spike width (Fig. 2c). DeepCAD-RT (1 ms: 0.190, 9 ms: 0.984) and PMD (1 ms: 0.554, 9 ms: 0.983) showed enhancement only when the spike width was larger than 5 ms and 3 ms, respectively, which verifies the importance of exploiting spatially adjacent pixels in the same time frame. We note that this inconsistency between the two metrics stems from the fact that the Pearson correlation coefficient is affected only by the time-varying component of the signals, whereas PSNR is largely determined by the static component.

For further comparison, we analyzed the voltage traces at the single pixel (Fig. 2d) and single cell levels (Fig. 2e). Only the single pixel voltage traces from SUPPORT retained the spike waveforms (Fig. 2d), whereas the spikes were buried under the noise level in that from the noisy video. DeepCAD-RT and PMD reduced the variance in the single pixel voltage traces, but the spikes were still not detectable due to the bias introduced by their signal models. The single cell voltage traces showed similar results (Fig. 2e), although the difference was less dramatic than the single pixel traces, as the SNR was improved by averaging multiple pixel values. SUPPORT was able to reduce variance without distorting the waveforms for every spike width. In comparison, the spikes were not detectable in the results from DeepCAD-RT and PMD when the spike width was under 3 ms. It should be noted that the performance of both DeepCAD-RT and PMD was better for larger spike widths, but for different reasons. DeepCAD-RT estimates the current frame given temporally adjacent frames, so the prediction becomes more accurate when the dynamics are slower. PMD attempts to find a low rank approximation of a given matrix that is supposedly closer to the ground truth, so a temporally long event is less likely to be “ignored” as its contribution to the approximation error is higher.

### Denoising single-neuron voltage imaging data

To validate SUPPORT on experimentally obtained data, we applied SUPPORT to *in vivo* single-neuron voltage imaging data with simultaneous electrophysiological recordings. The dataset contained light-sheet microscopy images of a single neuron in the dorsal part of the cerebellum of a zebrafish expressing Voltron1 with simultaneous cell-attached extracellular electrophysiological recording. Signals from electrophysiological recordings were recorded at 6 kHz, and light-sheet imaging was performed with a frame rate of 300 Hz^23^.

In the raw data, both the spatial footprint and temporal traces of the neuron were severely corrupted by Poisson– Gaussian noise. We compared temporal traces extracted from the raw video and the denoised video using SUPPORT, DeepCAD-RT, and PMD, along with the electrophysiological recording. Spike locations from the electrophysiological recordings were extracted (Methods) and visualized as black dots for a visual aid (Fig. 3a, b). After denoising with SUPPORT, the temporal trace showed a much lower variance compared to the temporal trace of the raw data while preserving the spikes. In comparison, while the temporal variance in the denoising outcome acquired using DeepCAD-RT was low, the spikes were no longer visible in the traces, which implies that the signal modeling in DeepCAD-RT significantly increased the bias. It should be noted that the subthreshold activity was recovered well in the outcomes of both SUPPORT and DeepCAD-RT, as temporally adjacent frames provide enough information to predict such low frequency components. The temporal trace from PMD was nearly identical to that from the raw video, which indicates that PMD had limited impact on both bias and variance.

**Fig. 3:**
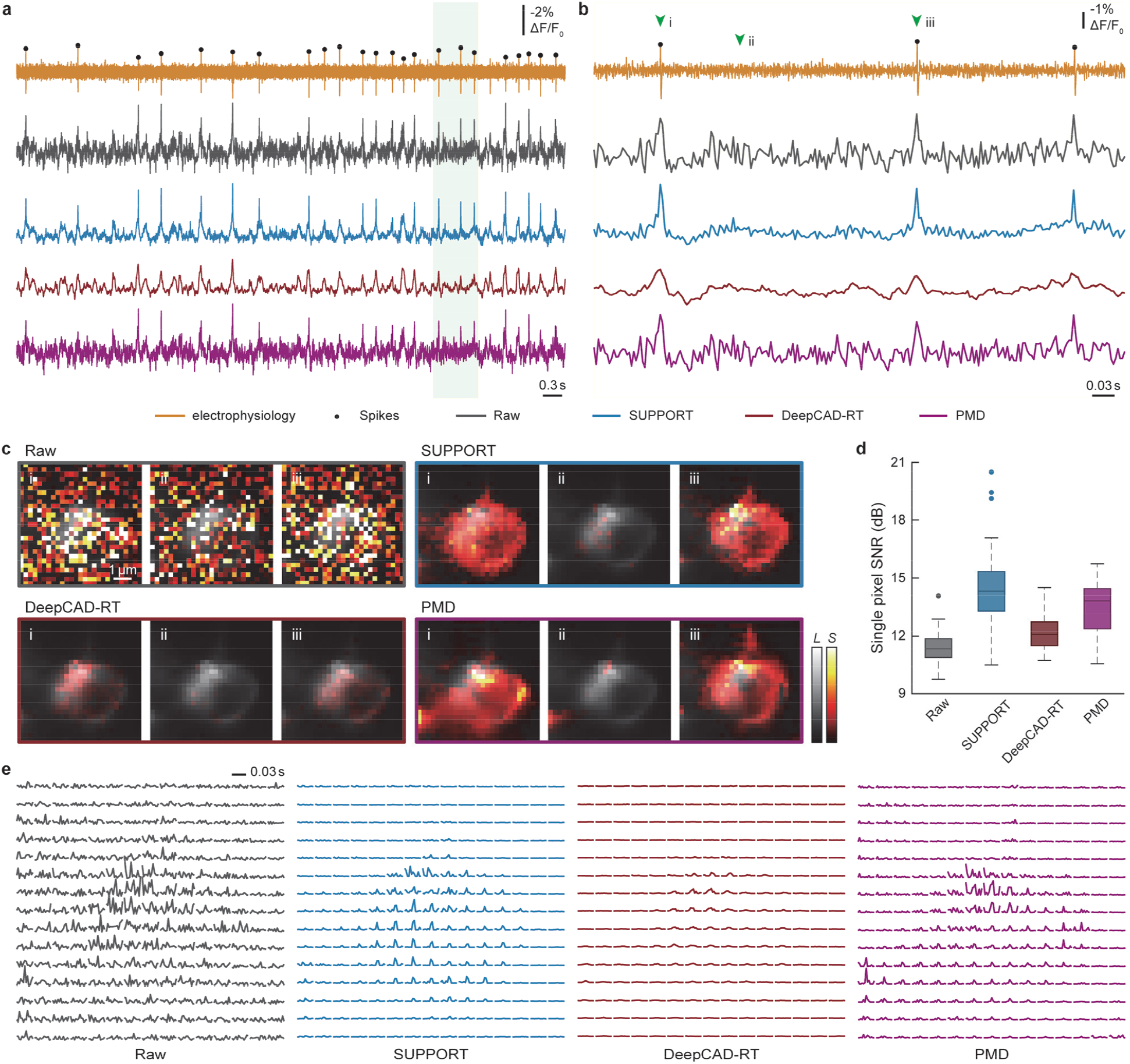
Denoising single-neuron voltage imaging data. **a**, Simultaneous electrophysiological recording and voltage imaging data. From top to bottom: Electrophysiological recording, raw, SUPPORT, DeepCAD-RT, and PMD denoised data. Detected spikes from electrophysiological recordings are marked with black dots. Traces from voltage imaging data were extracted using a manually drawn region of interest. **b**, Enlarged view of the green region in **a. c**, Three representative frames indicated on **b** with green arrows for raw and denoised data. Baseline and activity components are decomposed from raw data and denoised data. The baseline component with a gray colormap and the activity component with a hot colormap are overlaid. **d**, Box-and-whisker plot showing the signal-to-noise ratio for the pixels inside the cell region from raw and denoised data, *N* = 70. From left to right: Raw, SUPPORT, DeepCAD-RT, and PMD denoised data. **e**, Spatiotemporal diagram showing the voltage transients of each 2 × 2 binned pixel with a small temporal region centered at time point i on **b**. From left to right: Raw, SUPPORT, DeepCAD-RT, and PMD denoised data.

After we applied SUPPORT to enhance these data, not only did the neuronal activity become clearly visible in the images, but the spatial footprints of the activity also showed high consistency with the corresponding neuronal shape (Fig. 3c, Supplementary Video 2). Representative frames from the raw and denoised data show that SUPPORT removed the noise very effectively, while the activity was preserved.

For further comparison, we extracted single pixel fluorescence from the cell membrane pixels and found that the average single pixel SNR was strongly enhanced with SUPPORT (14.46 dB) compared to DeepCAD-RT (12.21 dB) and PMD (13.46 dB) (Fig. 3d). The spatiotemporal diagram, which visualizes the voltage transients of each 2 × 2 binned pixel, also verified that SUPPORT successfully reduced the variance while preserving the spikes at the pixel level (Fig. 3e). Additionally, we found that SUPPORT precisely revealed the traces from single pixels inside the soma (Supplementary Fig. 3) and along the axonal branch (Supplementary Fig. 4), which indicates SUPPORT’s suitability for studies involving voltage dependency along the axonal branch^34^. Finally, SUPPORT was able to denoise *in vitro* cultured neurons labeled with a synthetic voltage dye, which indicates its suitability for designing voltage indicators (Supplementary Fig. 5).

### Denoising population voltage imaging data

We applied SUPPORT to voltage imaging data that contained *in vivo* population neuronal activity in awake mouse cortex layer 1 expressing Voltron1^23^ and zebrafish spinal cord expressing zArchon1^27^. The mouse dataset was recorded with a wide-field fluorescence microscope with a frame rate of 400 Hz, and the zebrafish dataset was recorded with a light-sheet fluorescence microscope with a frame rate of 1 kHz^35^.

After applying SUPPORT to the voltage imaging data, we applied baseline correction (Methods). Despite the high noise level of the voltage imaging data, the neuronal structures became clearly visible after denoising (Fig. 4a, d, Supplementary Video 3). The single pixel SNR was improved by 9.11 dB on average (21.58 ± 1.62 dB for SUPPORT, 12.47 ± 0.89 dB for the raw data) for the mouse dataset (Fig. 4b), and 6.32 dB (19.08 ± 2.07 dB for SUPPORT, 12.72 ± 0.67 dB for the raw data) for the zebrafish dataset (Fig. 4e). For further analysis, we extracted the voltage traces from manually drawn ROIs (Fig. 4c, f, g). In line with the results from the simulation and the single-neuron voltage imaging, the variance was significantly decreased, while the sharp voltage transients induced by spikes were preserved.

**Fig. 4:**
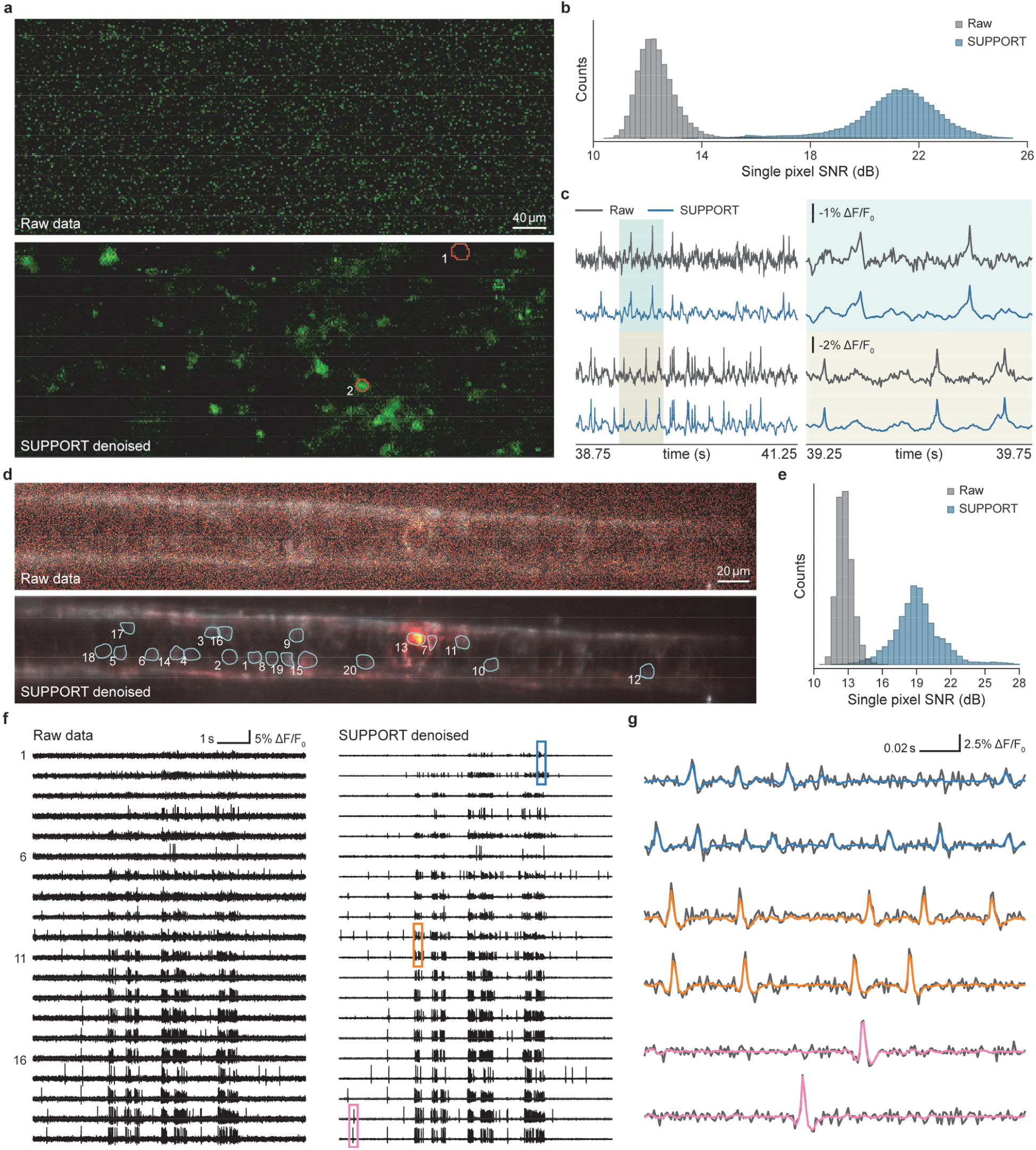
Denoising population voltage imaging data. **a**, Images after baseline correction from mouse dataset. Top: Baseline-corrected raw data. Bottom: Baseline-corrected SUPPORT denoised data (Methods). Boundaries of two regions of interest (ROIs) are drawn with red lines. **b**, Distribution of the signal-to-noise ratio (SNR) for all pixels from raw and SUPPORT denoised data after baseline correction, *N* = 65,536. **c**, Traces from raw and SUPPORT denoised data extracted from two ROIs in **a**. Traces for the smaller temporal region are plotted on the right. The enlarged temporal region is colored blue and brown. **d**, Images from the zebrafish dataset. Baseline and activity components are decomposed from raw data and SUPPORT denoised data (Methods). The baseline component with a gray colormap and the activity component with a hot colormap are overlaid. Boundaries of 20 ROIs are drawn with cyan lines. Top: Raw data. Bottom: SUPPORT denoised data. **e**, Distribution of SNR for pixels inside the ROIs from raw and SUPPORT denoised data, *N* = 5,722. **f**, Traces for 20 ROIs from raw and SUPPORT denoised data. Left: Raw data. Right: SUPPORT denoised data. **g**, Enlarged view of traces from colored regions in **f** are plotted. Traces from raw data are overlaid with a gray color and denoised data are overlaid with corresponding color in **f**.

We also extracted the neurons and corresponding temporal signals using localNMF^35^, which is an automated cell extraction algorithm, from the mouse and zebrafish datasets (Methods, Supplementary Fig. 6a, b). Owing to the improvement in SNR, we were able to automatically segment 42 neurons from the denoised mouse data compared to 31 neurons from the raw data. For zebrafish data, 27 neurons from the denoised data and 9 neurons from the raw data were extracted. We then measured the F_1_ score between the ground-truth ROIs and the extracted ROIs across several intersection-over-union (IoU) threshold values. We quantified the area under F_1_ score across the IoU curve, and there was a 1.6-fold improvement for mouse data (0.31 for denoised and 0.19 for raw data) and a 2.0-fold improvement for zebrafish data (0.43 for denoised and 0.21 for raw data) (Supplementary Fig. 6c). The extracted neuronal signal from SUPPORT also clearly shows spikes, while the signal from the raw data shows high variance (Supplementary Fig. 6d), which indicates that SUPPORT facilitates the automated analysis of large-scale population voltage imaging data.

It was shown that SUPPORT could denoise other population voltage imaging data with different regions and voltage indicators, indicating its suitability for the routine use of population voltage recordings (Supplementary Fig. 7, 8). Finally, we observed that SUPPORT trained on single population voltage imaging data accurately denoised another population voltage imaging data without fine-tuning (Supplementary Fig. 9), which demonstrates its generalizability.

### SUPPORT denoises freely moving *Caenorhabditis elegans* imaging data without bias

We tested SUPPORT for denoising three-dimensional time-lapse fluorescence microscopy images of *C. elegans*^36^, in which the differences among the frames came from the motion of the worm, which was not sampled with a sufficiently high imaging speed. The nuclei of all neurons in the worm were labeled using red fluorescent protein mCherry^37^ under the H20 promoter. The volume images with 20 axial slices were recorded with spinning disk confocal microscopy at a volume rate of 4.75 Hz.

We denoised the video using SUPPORT, DeepCAD-RT, and PMD in a plane-by-plane manner. We first compared the noisy data and the denoised results for a single axial slice. SUPPORT successfully denoised the images without any visible artifacts, whereas the denoising outcomes acquired using DeepCAD-RT and PMD suffered from motion-induced artifacts (Fig. 5a), which again proves the importance of employing an appropriate signal model for denoising. The difference between the SUPPORT output and the noisy input, which was expected to be white noise, did appear purely white. However, the difference between the outputs from DeepCAD-RT and PMD and the noisy input contains low frequency components that are highly correlated with the structure of the input image (Fig. 5b).

**Fig. 5:**
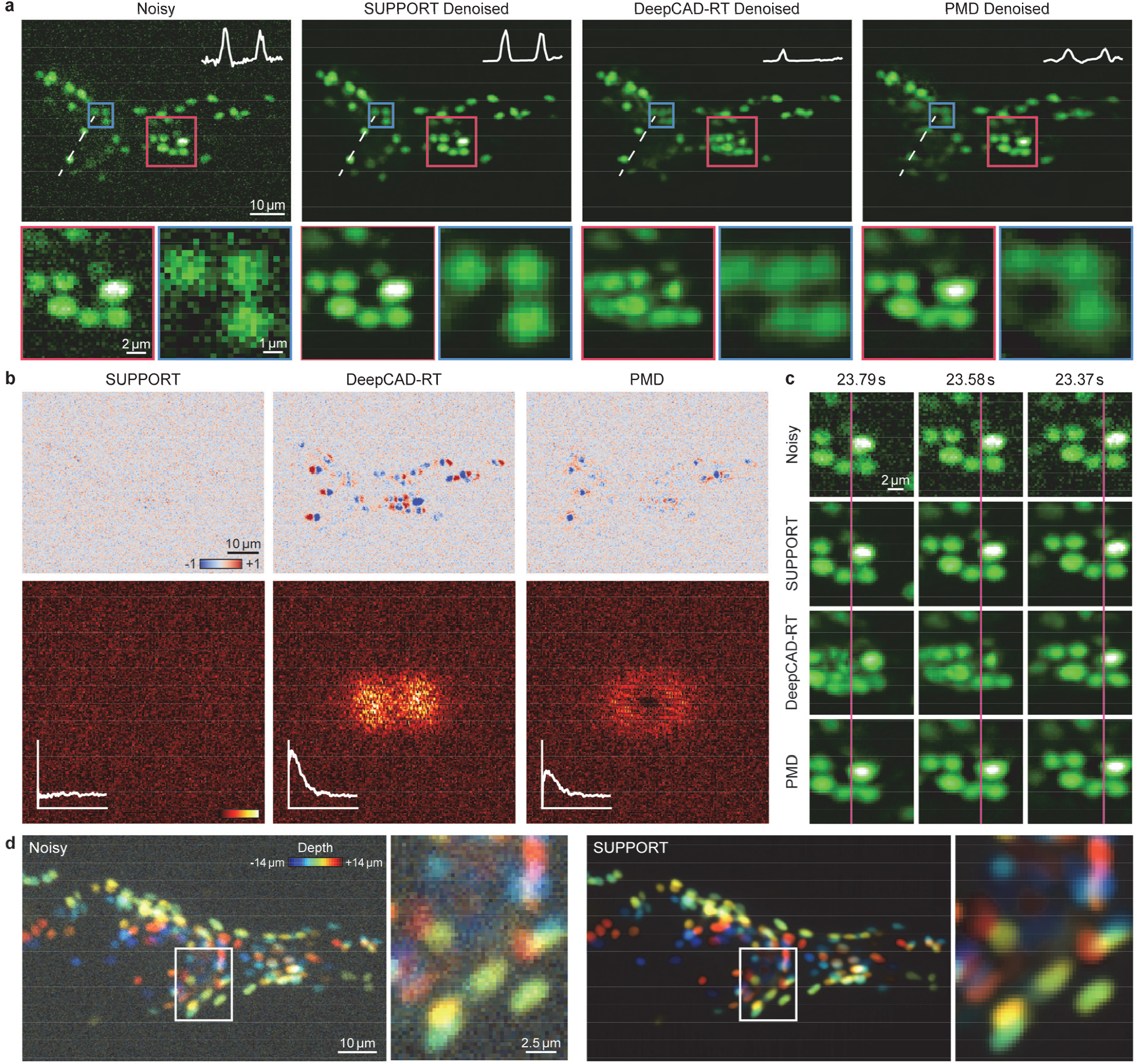
SUPPORT denoises freely moving *Caenorhabditis elegans* imaging data without bias. **a**, Images of freely moving *C. elegans*. From left to right: Noisy, SUPPORT, DeepCAD-RT, and PMD denoised data. Inset shows the intensity profile along the dashed line. Magnified views of the boxed regions are presented underneath. **b**, Pixel-wise difference between denoised data and noisy data. Squared norm of Fourier transform of each difference are shown in the lower images. Inset shows the logarithm of the squared norm of Fourier transform against the distance to the origin. **c**, Magnified views of the red boxed region in **a** at consecutive neighboring time points. Magenta lines were set on the left side of the brightest neuron in the noisy data. From top to bottom: Noisy, SUPPORT, DeepCAD-RT, and PMD denoised data. **d**, Noisy volume and denoised volume are depth coded and presented. Magnified views of the boxed regions are presented on the right.

In the consecutive frames shown in Fig. 5c, the worm’s locomotion is considerably faster than the imaging speed, which precludes the accurate prediction of the current frame based on adjacent frames. Nevertheless, SUPPORT successfully denoised the image without suffering from motion artifacts by incorporating information from neighboring pixels in the current frame, whereas DeepCAD-RT and PMD failed to predict the location of each cell, which was manifested as motion-induced artifacts in the images. The volumetric denoising outcome (Fig. 5d, Supplementary Video 4) demonstrates that SUPPORT can be used for denoising not only functional imaging data but also volumetric time-lapse images in which the speed of dynamics is faster than the imaging speed.

### SUPPORT denoises volumetric structural imaging data

To demonstrate the generality of SUPPORT, we evaluated it on denoising volumetric structural imaging data in which no temporal redundancy could be exploited for denoising. SUPPORT was tested on two volumetric datasets that contained *Penicillium* imaged with confocal microscopy and mouse embryos imaged with expansion microscopy^38^. *Penicillium* was imaged with two different recording settings to generate a pair of low-SNR and high-SNR volumes (Methods).

The volumetric images were denoised with SUPPORT regarding each z-stack as a time series. The qualitative analysis showed that SUPPORT was able to enhance the signal of volumetric structural imaging data, revealing the structures that were hidden by the noise (Fig. 6a, b, e, f, Supplementary Fig. 10, Supplementary Video 5). The fine structure of *Penicillium* was recovered with SUPPORT (Fig. 6d), which shows that its signal model is capable of learning statistics from a wide range of data. For the quantitative evaluation of SUPPORT with the *Penicillium* dataset, the Pearson correlation coefficients and SNR were measured by regarding the high-SNR image as a ground truth for each plane along the z-axis (Fig. 6c). The average Pearson correlation coefficients of SUPPORT (0.7625 ± 0.0742) showed 0.2960 increments compared to the low-SNR image (0.4665 ± 0.0904) and the average SNR of SUPPORT (8.6480 ± 0.6218 dB) showed 5.9792 dB increments compared to the low-SNR image (2.6688 ± 0.5130 dB). The qualitative and quantitative studies showed that SUPPORT is capable of enhancing not only time-lapse images but also static volumetric images. Thus, SUPPORT can be used in a wide range of biological research incorporating microscopic imaging.

**Fig. 6:**
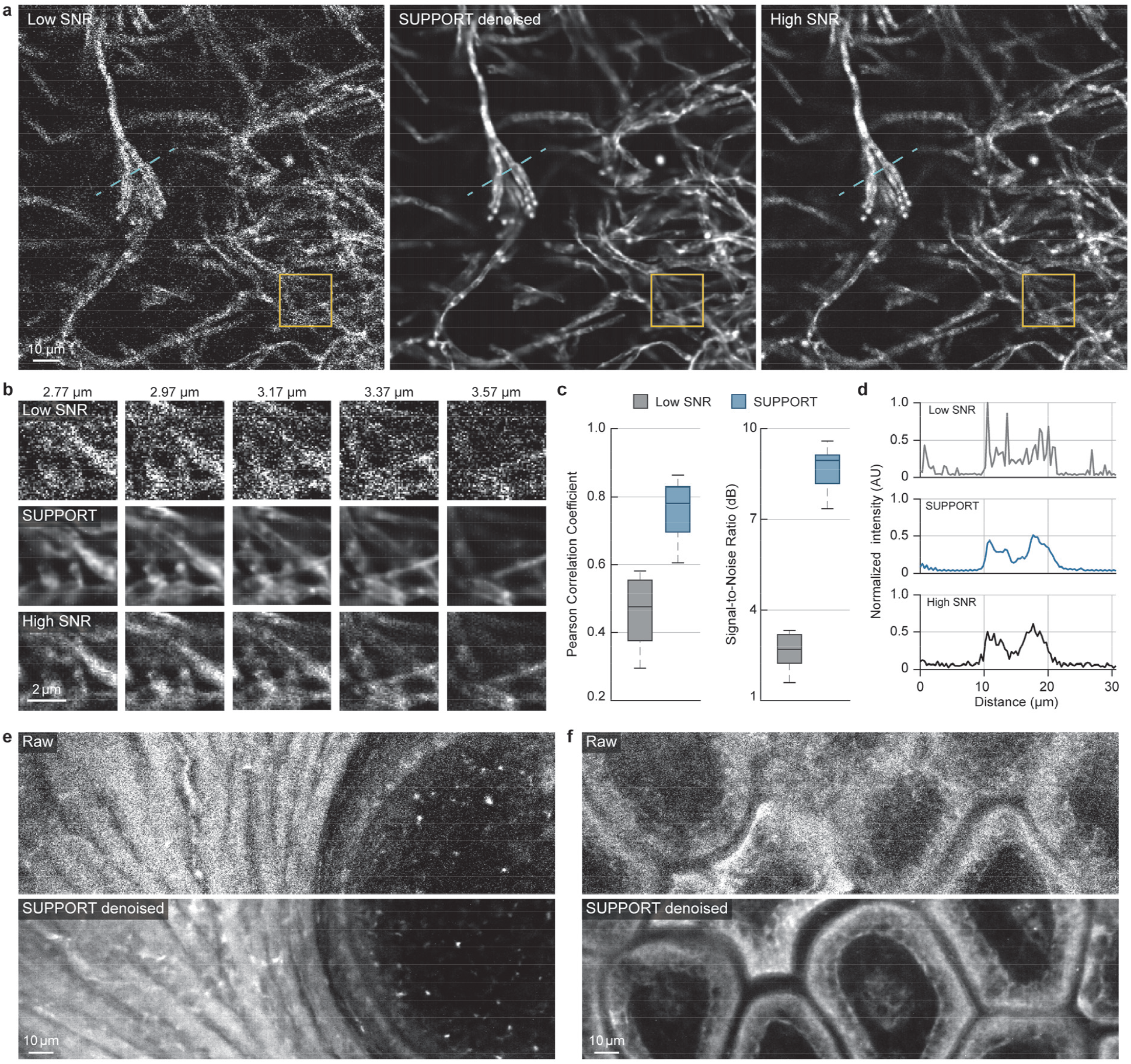
SUPPORT denoises volumetric structural imaging data. **a**, Representative axial slice from low-SNR, SUPPORT denoised, high-SNR volumes of *Penicillium*. **b**, Magnified views of the yellow boxed region in **a** at multiple axial locations. Axial location of **a** corresponds to 3.37 μm. **c**, Box-and-whisker plot showing Pearson correlation coefficient and signal-to-noise ratio for axial slices, *N* = 381. **d**, Intensity profiles of the cyan dashed line in **a. e**, Example frame of bone of a mouse embryo after expansion for the raw data (top) and denoised image using SUPPORT (bottom). **f**, Raw (top) and denoised image (bottom) of intestine of a mouse embryo. **e-f**, Length scales are presented in pre-expansion dimensions.

## DISCUSSION

SUPPORT is a self-supervised denoising method with a demonstrated ability to denoise diverse voltage imaging datasets acquired using Voltron1, Voltron2, paQuasAr3-s, QuasAr6a, zArchon1, and BeRST1. Thanks to its statistical prediction model that predicts a pixel value *x*_i,k_ by integrating the information from its spatiotemporal neighboring pixels Ω_i,k_ (i.e., 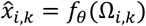), it showed high robustness when faced with the fast dynamics in the scene. While this design allows SUPPORT to simultaneously achieve low bias and low variance, it still leaves room for fundamental improvement, as it does not exploit the information contained in *y*_i,k_. The reason *y*_i,k_ was not exploited as an input is because it is used as the target in place of the ground truth for self-supervised learning; we cannot use *y*_i,k_ as both the input and the target of the network, as the network will simply become an identity function. This means that the cost of truly exploiting all available information is to give up the self-supervised learning scheme that does not require ground truth.

Denoising time-lapse imaging data in which a *C. elegans* exhibited rapid movement and a single volumetric image demonstrated that SUPPORT is not limited to denoising voltage imaging data; it can be used for denoising any form of time-lapse imaging data including calcium imaging (Supplementary Fig. 11–15, Supplementary Video 6) in which the imaging speed is slow compared to the underlying dynamics or volumetric structural imaging data. This is an important finding, as it indicates that the data do not need to be low rank to be denoised using SUPPORT, which is often required by many denoising algorithms^6,39^. Also, SUPPORT was able to be trained with only 3000 frames (Supplementary Fig. 16), which would facilitate its general usage in many laboratories with common desktop settings, especially with our GUI-based SUPPORT (Supplementary Fig. 17). Overall, its self-supervised learning scheme, robustness to fast dynamics, low variance in denoising outcomes, and compatibility with motion make it a versatile tool for processing a wide range of image data.

## METHODS

### SUPPORT network architecture and training details

The architecture of the SUPPORT network consists of two sub-networks: 2D U-Net and the blind spot network. 2D U-Net exploits the information of the temporally adjacent frames. The input data are first separated into two blocks: (1) temporal neighboring frames and (2) the center frame. The temporal neighboring frames are concatenated in the channel dimension and passed through 2D U-Net. Then, the center frame and the output of 2D U-Net are concatenated in the channel dimension and passed through the blind spot network, which has a zero at the center of the impulse response. Finally, the outputs of 2D U-Net and the blind spot network are concatenated in the channel dimension and passed through 1 × 1 convolution layers. The overall architecture is illustrated in Supplementary Fig. 1a.

2D U-Net^31^ consists of a 2D encoder, a 2D decoder, and skip connections from the encoder to the decoder (Supplementary Fig. 1b). In the 2D encoder, there are four encoder blocks. Each block consists of a 3(*x*) × 3(*y*) convolutional layer, followed by a BatchNorm, a LeakyReLU, and a 2(*x*) × 2(*y*) max pooling layer. In the decoder, there are four decoder blocks, each of which contains a bilinear interpolation followed by a 3(*x*) × 3(*y*) convolutional layer, a BatchNorm, and a LeakyReLU. The skip connections link low- and high-level features by concatenating feature maps in the channel dimension. We designed 2D U-Net to take the previous 30 frames and next 30 frames as the input. For denoising structural imaging data, the previous 10 frames and next 10 frames were used as the input.

The blind spot network was designed to efficiently increase the receptive field of the network over computation (i.e., memory and the number of multiply-add operations). A comparison between previous blind spot network designs^19,30^ is shown in Supplementary Fig. 2. The blind spot network consists of 1) two sequential parts and 2) an aggregating part (Supplementary Fig. 1c). There are two sequential paths that use convolutional layers with kernel sizes of 3 × 3 and 5 × 5. Each sequential path consists of sequential blind spot convolutional layers with “shortcut connections” (Supplementary Fig. 1c). The center value of the weight of the blind spot convolutional layer is masked as 0 to make the blind spot property. For the kernel size of 3 × 3, the dilation and padding are both set as 2^i^ for the *i*-th layer to preserve blind spot properties for each feature after the layer. Similarly, for the kernel size of 5 × 5, the padding and dilation are set as 2 × 3^i^. The shortcut connection links the input to the features by adding the input, passed by the 1 × 1 convolutional layer, to the intermediate features. In the aggregating path, all features after each layer in the sequential paths are concatenated in the channel dimension and then passed through three 1 × 1 convolutional layers to finally predict the signal. The receptive field of the blind spot network is illustrated in Fig. 1b, which shows the fractal-like pattern.

For the data in which structured noise can be predicted from the neighboring pixels, options to change the size of the blind spot were also implemented (Supplementary Fig. 11). To increase the size of the blind spot to *p*, we added additional dilation and padding of ⌊*p*/2⌋ for the last blind spot convolutional layers of two sequential paths. Also, only the final features of two sequential paths, rather than all intermediate features, were passed through the aggregating path.

The network was trained on Pytorch 1.12.1 and CUDA 11.3 with an NVIDIA RTX 3090 GPU and an Intel Xeon Silver 4212R CPU. For the loss function, the arithmetic average of L1-loss and L2-loss were used. As a preprocessing step, each input video was normalized by subtracting the average value and dividing by the standard deviation. For data augmentation, random flipping and rotation by integer multiples of 90 degrees were used. An Adam optimizer^40^ with a learning rate of 5 × 10^−4^ was used for gradient-based optimization. Training SUPPORT for processing the zebrafish dataset which had a size of 1,024 (*x*) × 148 (*y*) × 24,000 (*t*) took 47 hours for 14 million gradient updates. The inference for the same dataset took 30 minutes.

### Synthetic voltage imaging data generation

Simulating synthetic voltage imaging data includes the pipeline of first generating clean video (ground truth) and then adding Poisson and Gaussian noise. To generate a realistic spatial profile that resembles neurons in a mouse brain, we used a NAOMi^32^ simulator that was originally developed for simulating a two-photon calcium imaging dataset. The code was modified to generate voltage transients instead of calcium transients as temporal components. For the temporal components that define the fluorescence fluctuations of each neuron, we generated five different temporal components with a total of 15,000 frames and a frame rate of 500 Hz with different spike widths, ranging from 1 ms to 9 ms. The constructed voltage signals were matched to the parameters of Voltron, including spike frequency, *dF/F*_*0*_, and the level of subthreshold activity. Every other parameter was set as default apart from increasing the simulated field of view (FOV) two-fold. The noisy video was generated by adding Poisson and Gaussian noise. To add Poisson noise to the images, we first normalized the input images and multiplied them by 1000, and then used each pixel value as the parameter (i.e., mean value) of the Poisson distribution. Thereafter, Gaussian noise with a mean of 0 and a standard deviation of 5 was added to the images. Finally, negative values were truncated to 0.

### L1 and L2 loss comparison with simulation and real data

The dependency of the denoising performance and loss function was investigated through denoising simulation and real data. The weighted average of L1 and L2 loss, *ℒ* = α*ℒ*_1_ + (1-α)*ℒ*_2_, with α {0, 0.3, 0.5, 0.7, 1} for simulated data and α {0, 0.5, 1} for real data were used as a loss function. For simulation, synthetic voltage imaging data generated using a NAOMi simulator with a spike width of 3 ms were used. For real data, simultaneous electrophysiological data with voltage imaging were used. The data were fed into the network training procedure with different loss functions (Supplementary Fig. 18, 19).

### In vivo single-neuron simultaneous voltage recording and electrophysiology

The data from simultaneous light-sheet fluorescence imaging and cell-attached extracellular recordings of single-neuron activity were acquired from the study of Voltron1^23^. A recording of the dorsal part of the cerebellum of a zebrafish expressing Voltron1 under control of GAL4-UAS system and *vGlut2a* promoter (*Tg(vGlut2a:GAL4);Tg(UAS:Voltron1)*) was used in this study.

The data from simultaneous structured illumination fluorescence imaging and patch-clamp electrophysiological recordings of single-neuron activity were recorded with mouse cortex L2/3 pyramidal neurons using a DMD with a frame rate of 1000 Hz. Voltron2 and QuasAr6a were expressed using in-utero electroporation.

An ROI was manually drawn around the neuron, and fluorescence traces were extracted from the mean signal of the ROI in the temporal stack. The baseline estimation of fluorescence was performed to account for photobleaching by calculating the 80th quantile. The estimated baseline was used to calculate Δ*F*/*F*_O_. Action potentials from fluorescence traces and electrophysiological recordings were inferred by thresholding.

### In vitro single-neuron voltage recording

We prepared primary rat hippocampal neurons cultured on a 35 mm glass bottom dish (P35G-1.5-14-C, MatTek). At nine days in vitro, neurons were stained with a voltage-sensitive dye (BeRST1, 2 μM) dissolved in an imaging solution containing 140 mM NaCl, 3 mM KCl, 3 mM CaCl_2_, 1 mM MgCl_2_, 10 mM HEPES, and 30 mM glucose (pH 7.3) for 15 min, and then rinsed with a fresh imaging solution prior to optical imaging^28^. Time-lapse imaging of spontaneous neural activity was acquired using an inverted microscope (Eclipse Ti2, Nikon) equipped with a 40× water immersion objective lens (NA 1.15; MRD7710, Nikon), while maintaining the sample temperature at 30°C. For excitation, an LED (SOLIS-623C, Thorlabs) with a bandpass filter (ET630/20x, Chroma Technology) was used at an irradiance of 20 mW/mm^2^ at the sample. Emission was passed through a dichroic mirror (T660lpxr, Chroma Technology) and an emission filter (ET665lp, Chroma Technology), and was collected by an sCMOS camera (Orca Flash 4.0, Hamamatsu) at a 1-kHz frame rate with 4 × 4 binning and subarray readout (361 × 28 pixels) for a duration of 25 s.

### In vivo single-neuron simultaneous calcium recording and electrophysiology

A craniotomy over V1 was performed, and neurons were infected with adeno-associated virus (AAV2/1-hSynapsin-1) encoding jGCaMP8f. At 18–80 days after the virus injection, the mouse was anesthetized, the cranial window was surgically removed, and a durotomy was performed. The craniotomy was filled with 10–15 μL of 1.5% agarose, and a D-shaped coverslip was secured on top to suppress brain motion and leave access to the brain on the lateral side of the craniotomy. The mice were then lightly anesthetized and mounted under a custom two-photon microscope. Two-photon imaging (122 Hz) was performed of L2/3 somata and neuropil combined with a loose-seal, cell-attached electrophysiological recording of a single neuron in the field of view. Temporally fourfold downsampling was held to the data to reduce the sampling rate before the analysis. After excluding some outlier recordings with a low correlation between calcium signal and action potentials, an ROI was manually drawn around the neuron, and fluorescence traces were extracted from the mean signal of the ROI in the temporal stack.

### Neuronal population recordings

Voltage and calcium imaging data of large neuronal populations with different brain regions, cell types, recording rates, reporters, and imaging modalities were used to validate the general applicability of SUPPORT. The properties of the datasets are reported in Supplementary Table 1.

### Volumetric structural imaging of *Penicillium*

For the volumetric structural imaging of *Penicillium*, the specimen was imaged using a point-scanning confocal microscopy system (C2 Plus, Nikon) equipped with a 16× 0.8 NA water dipping objective lens (CFI75 LWD 16X W, Nikon). The imaging was performed using a 488 nm excitation laser with a laser power of 0.075 mW for the low-SNR image and a laser power of 1.5 mW for the high-SNR image. The frame rate was 0.5 Hz for 1024 × 1024 pixels with a pixel size of 0.34 μm and each volume consisted of 1000 z-slices with a z-step size of 0.1 μm.

### Volumetric structural imaging of mouse embryos using expansion microscopy

Mouse embryos were isolated on day 15.5 of pregnancy in C57BL/6J mice and fixed with ice-cold fixative (4% paraformaldehyde in 1× phosphate buffered saline) for a day at 4°C. Fixed mouse embryos were embedded in 6% (w/w) low-gelling-temperature agarose and then sliced to a thickness of 500 μm with a vibratome. Embryo slices were then processed for anchoring, gelation, Alexa Flour 488 NHS-ester staining, digestion, decalcification, and expansion according to the previously described whole-body ExM protocol^38^. Following a 4.1-fold expansion of the embryo slices in the hydrogel, the sample was attached to cover glass and imaged using a confocal microscope (Nikon Eclipse Ti2-E, Tokyo, Japan) with a spinning disc confocal microscope (Dragonfly 200; Andor, Oxford Instruments, Abingdon, UK) equipped with a Zyla 4.2 sCMOS camera (Andor, Oxford Instruments) and a 10× 0.45 NA air lens (Plan Apo Lambda, Nikon). Z-stack images were obtained with a z-step size of 1 μm for intestine and bone, and 0.5 μm for tail. All animal experiments involving mouse embryos conducted for this study were approved by the Institutional Animal Care and Use Committee (IACUC) of Korea Advanced Institute of Science and Technology (KA2021-040).

### Confocal imaging of zebrafish neuronal calcium signals

For zebrafish experiments, transgenic larval zebrafish (*Danio rerio*) expressing GCaMP7a calcium indicator under control of GAL4-UAS system and *huc* promoter (*Tg(huc:GAL4);Tg(UAS:GCaMP7a)*)^41–43^ with a *Casper* (*mitfa(w2/w2);mpv17(a9/a9)*)^44^ mutant were imaged at 3–4 days post-fertilization (dpf). Zebrafish were maintained under standard conditions at 28°C and a 14:10 hour light:dark cycle.

The larvae were paralyzed by bath incubation with 0.25 mg/ml of pancuronium bromide (Sigma-Aldrich) solution for 2 mins^45^. After paralysis, the larvae were embedded in agar using a 2% low melting point agarose (TopVision) in a Petri dish. The dish was filled with standard fish water after solidifying the agarose gel. Specimens were imaged using a point-scanning confocal microscopy system (C2 Plus, Nikon) equipped with a 16× 0.8NA water dipping objective lens (CFI75 LWD 16X W, Nikon). The imaging was performed using a 488 nm excitation laser (0.15–0.75 mW) with a frame rate of 1–4 Hz and a pixel size of 0.34–0.75 μm. Multiple brain regions were imaged at the frame rate of 1 Hz for 1024 × 512 pixels with a pixel size of 0.75μm. The cerebellar plate, dorsal telencephalon, medulla oblongata, olfactory bulb, and optic tectum were imaged at a frame rate of 2 Hz for 1024 × 256 pixels with a pixel size of 0.34 μm, and the habenula was imaged at a frame rate of 4 Hz for 512 × 128 pixels with a pixel size of 0.34 μm. All animal experiments involving zebrafish conducted for this study were approved by the Institutional Animal Care and Use Committee (IACUC) of Korea Advanced Institute of Science and Technology (KA2021-125).

### Baseline and activity decomposition of voltage imaging data

For visualization, the data were decomposed into the underlying baseline and neuronal activity. The baseline estimation was performed using the temporal moving average. Window length was chosen in accordance with the recording rate for the data. For the data that only required photobleaching correction, b-spline fit was used to estimate baseline without using the moving average (Fig. 4a). For the positive-going voltage indicators (zArchon1, QuasAr6a, paQuasAr3-s), the activity component was acquired by subtracting the estimated baseline from the data. For the negative-going voltage indicators (Voltron1, Voltron2), the activity component was acquired by subtracting the data from the estimated baseline.

### Cell detection of neuronal populations of voltage and calcium imaging data

For voltage imaging data, the SGPMD-NMF pipeline was applied to detect ROIs and corresponding temporal signals, which is available on GitHub (https://github.com/adamcohenlab/invivo-imaging). In the pipeline, detrending based on b-spline fitting and demixing based on localNMF were used without additional denoising. For the mouse cortex data, detrended data was flipped before the demixing step by subtracting the data from the maximum value of the data, since the data were recorded with Voltron1, which is a negative-going voltage indicator. After extraction, we removed non-neuronal spatial components with the following simple heuristics:

1. Reject if the number of pixels in the component is smaller than α.
2. Reject if the width or height of the component is larger than β.
3. Reject if the *width*/*height* is not in [γ, δ].

For mouse cortex data, only the first heuristic was used, with α set as 10, where the size of the neurons was small in the data. For zebrafish data, all heuristics were used, with α = 100, β = 50, γ = 0.5, and, δ = 1.5 (Supplementary Fig. 6).

Cellpose^46^ was applied to a single frame image of SUPPORT denoised video to detect cells (Supplementary Fig. 15). All parameters were set to default except ‘flow’ and ‘cellprob’, which were set by empirical values that best fit the data.

### Performance metrics

Signal-to-noise ratio (SNR), peak signal-to-noise ratio (PSNR), and root-mean-squared error (RMSE) were used as metrics to evaluate the pixel-level consistency between SUPPORT denoised images and ground-truth images. The RMSE between the signal *x* and the reference signal *y* is defined as 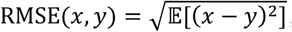, where 𝔼 denotes the arithmetic mean. The SNR between the signal *x* and the reference signal *y* is defined as 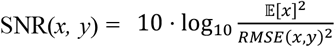. The PSNR between the signal *x* and the reference signal *y* is defined as 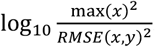. Pearson’s ate the consistency between denoised traces and ground-truth traces. The Pearson correlation between the signal *x* and the reference signal *y* is defined as 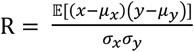 where *μ*_x_ and *μ*_y_ are the mean values of signal *x* and *y*, respectively, and *σ*_x_ and *σ*_y_ are the standard deviations of signal *x* and y, respectively. As a measure of the performance of denoising experimental voltage imaging data, SNR was used with modification, as the ground truth was not available. The SNR of the signal *x* is defined as 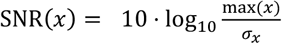. log ^max(x)^. Metrics were calculated after baseline correction.

For the population voltage imaging data, we measured the performance of cell extraction. After cell detection using localNMF, we first calculated IoU between all pairs of extracted components and manually segmented cells. The cell extraction was given as correct if the IoU between the extracted component and the manual segmentation was higher than the threshold. Then, the F_1_ score (i.e., harmonic mean of precision and recall) was calculated.

### SUPPORT with graphical user interface

We developed SUPPORT software with a graphical user interface. Using this software, arbitrary functional imaging data can be denoised in two steps: (1) Loading: Load the data and pretrained SUPPORT model, and (2) Running: Denoise the data, visualize raw and denoised frames in real time, and save denoised data.

### Comparison with penalized matrix decomposition (PMD)

We used the publicly available implementation of PMD software from the official GitHub repository (https://github.com/ikinsella/trefide). All parameters were set to default except patch size, according to the pixel size of the raw data.

### Comparison with DeepCAD-RT

We used the publicly available Pytorch implementation of DeepCAD-RT from the official GitHub repository (https://github.com/cabooster/DeepCAD-RT). All parameters were set to default except patch size.

## ACKNOWLEDGEMENTS

The zebrafish lines used for calcium imaging were provided by the Zebrafish Center for Disease Modeling (ZCDM), Korea. This research was supported by National Research Foundation of Korea (NRF) (2020R1C1C1009869, 2021R1A4A102159411), Bio & Medical Technology Development Program through the NRF funded by the Ministry of Science and ICT (NRF-2021M3A9I4026318), Original Technology Program through the NRF funded by the Ministry of Science and ICT, Republic of Korea (2021M3F3A2A01037808), and the BK21 plus program through the NRF funded by the Ministry of Education of Korea.

## DATA AVAILABILITY

The dataset of light sheet imaging of Voltron1 expressing zebrafish spinal cord neurons can be downloaded from (https://figshare.com/articles/dataset/Voltage_imaging_in_zebrafish_spinal_cord_with_zArchon1/14153339). The dataset of light sheet imaging of Voltron1 expressing zebrafish tegmental area can be downloaded from (https://figshare.com/articles/dataset/Voltron_Behavior_imaging/9638756). The dataset of confocal imaging of mCherry expressing *C*.*elegans* can be downloaded from (https://figshare.com/articles/dataset/Materials_for_Accurate_automatic_detection_of_densely_distributed_cell_nuclei_in_3D_space_/3184546/1). The dataset of simultaneous two-photon imaging and electrophysiological recording of GCaMP6f expressing mouse cortex V1 neurons can be downloaded from the Collaborative Research in Computational Neuroscience (CRCNS) at (http://crcns.org/data-sets/methods/cai-1). The dataset of confocal imaging of zebrafish neuronal calcium signals and volumetric structural imaging of *Penicillium* has been made publicly available at (https://github.com/NICALab/SUPPORT).

## CODE AVAILABILITY

Code for Pytorch implementation of SUPPORT is available online at Github repository (https://github.com/NICALab/SUPPORT).

## AUTHOR CONTRIBUTIONS

M.E. and S.H. designed denoising algorithm. M.E. and S.H. designed and performed experiments and analyzed data.

G.K. performed calcium imaging data analysis. E.-S.C. performed confocal imaging of zebrafish and Penicillium. J.S. performed expansion microscopy imaging of mouse embryos under supervision of J.C. P.P. performed simultaneous electrophysiology and voltage imaging experiments under supervision of A.E.C. K.-H.L performed zebrafish experiments under supervision of C.-H.K. S.K. performed in vitro single-neuron voltage recording under supervision of M.C. M.R. performed in vivo single-neuron simultaneous calcium recording and electrophysiology under supervision of K.S. M.E., S.H., and Y.-G.Y. wrote the manuscript with input from all authors. Y.-G.Y. conceived and led this work.

## COMPETING FINANCIAL INTERESTS

The authors declare no competing financial interests.

## Notes

### Competing Interest Statement

The authors have declared no competing interest.

