## Supplementary information for "Statistically unbiased prediction enables accurate denoising of voltage imaging data"

**Supplementary Figures 1–19**

**Supplementary Videos 1–6**

**Supplementary Tables 1**

**References**

### SUPPLEMENTARY FIGURES

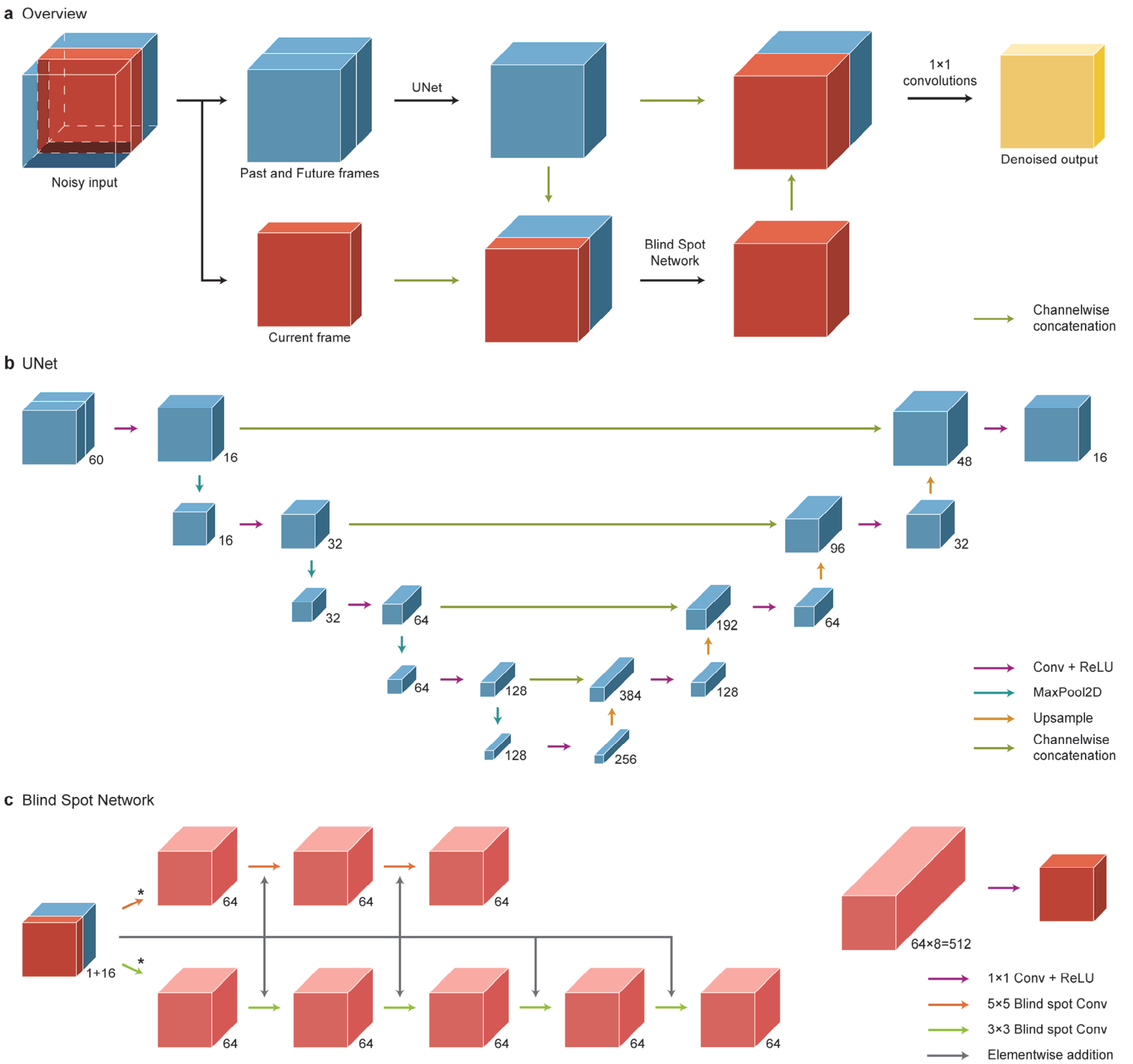


**Supplementary Fig. 1: Network architecture. a**, Overall architecture of the network. The network predicts the center value based on its spatiotemporally neighboring pixels. Pixels in past and future frames are processed by 2D U-Net, and spatially neighboring pixels in the current frame are processed by a blind spot network. **b**, 2D U-Net, which consists of a 2D encoder, a 2D decoder, and skip connections, processes the past and future frames. The past and future frames are channelwise concatenated and given to the 2D U-Net as input. **c**, The blind spot network consists of blind spot convolution layers, which are convolution layers with zero at the center of the kernels. In addition to the current frame, feature maps from the 2D U-Net, which encode the information in temporally neighboring frames, are channelwise concatenated and given as input to the blind spot network. (*) The first blind spot convolution layer has a blind spot property only in the first channel, which corresponds to the current frame. For the other layers, there are blind spots in all channels.

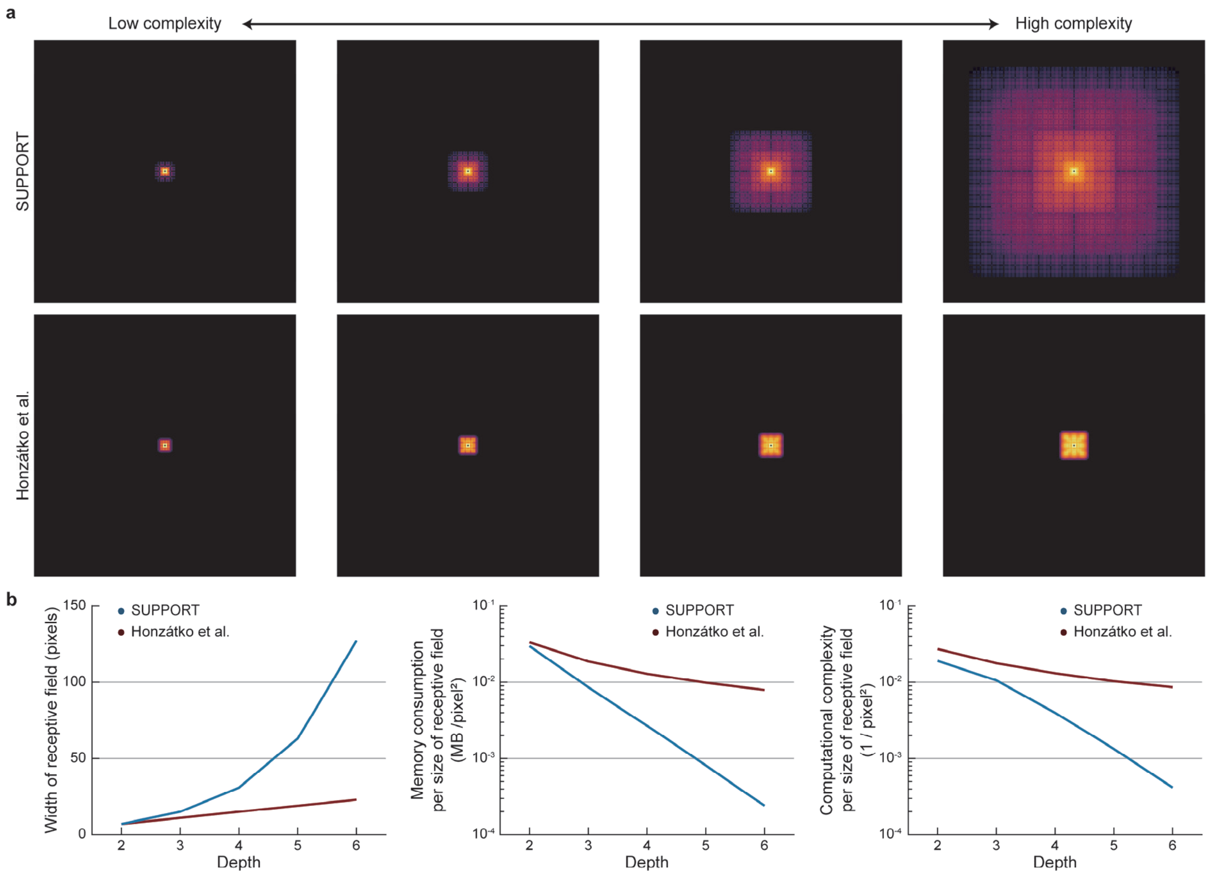


**Supplementary Fig. 2: Comparison of blind spot networks with different complexities. a**, Impulse response of two blind spot network architectures (SUPPORT, Honzátko et al.[1]) with different network complexities. While the width of SUPPORT's receptive field increases exponentially, Honzátko et al. increases linearly. **b**, The width of the receptive field, memory consumption, and the number of multiply-add operations over the size of the receptive field of SUPPORT and Honzátko et al. for different depths.

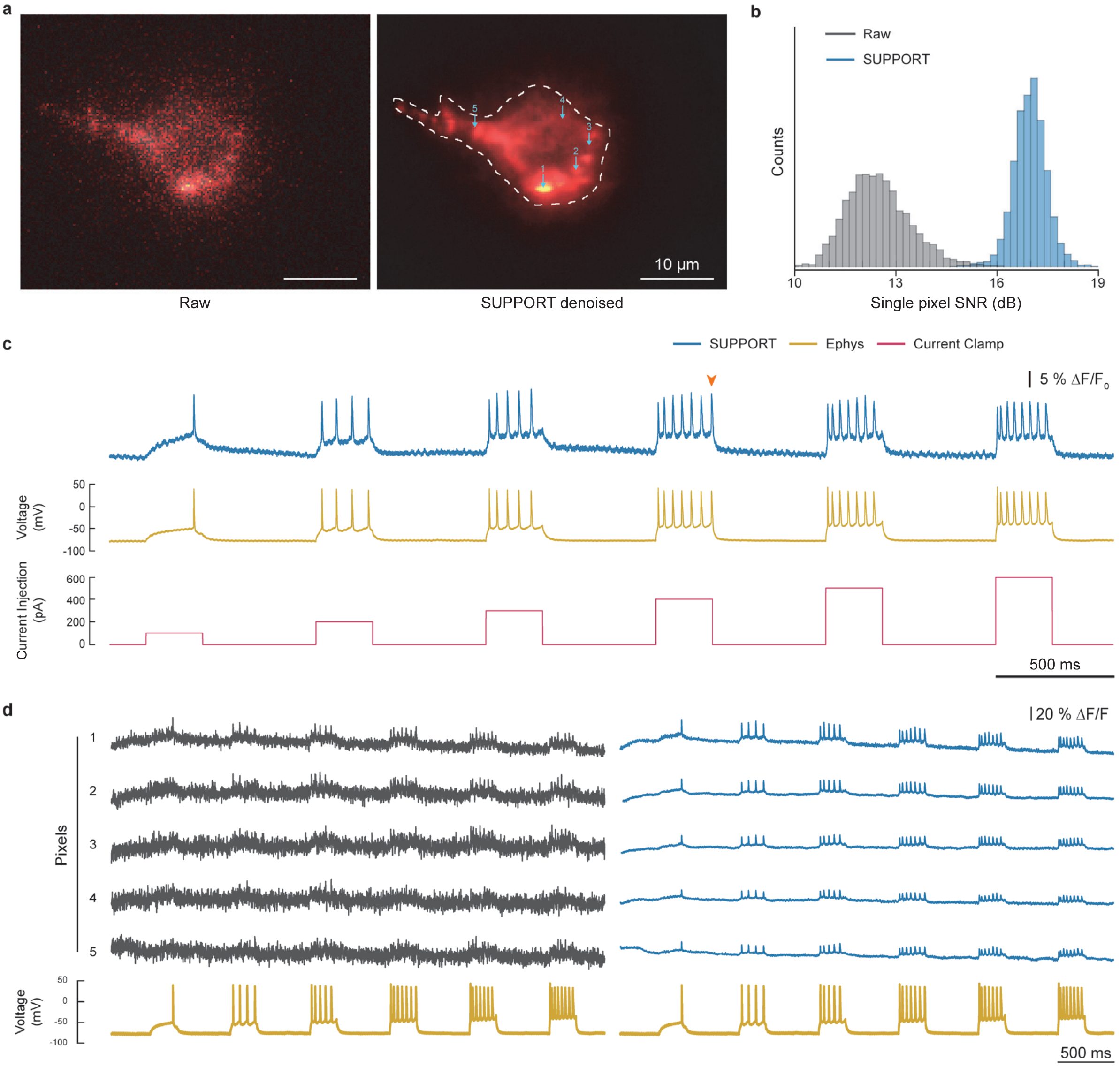


**Supplementary Fig. 3: SUPPORT reveals single pixel traces from voltage imaging data with simultaneous electrophysiological recording. a**, Representative frames of raw and SUPPORT denoised videos are shown after baseline correction. The frames used are indicated on **c** with an orange arrow. QuasAr6a-expressing mouse cortex L2/3 was used as a dataset (Supplementary Table 1). The baseline component with gray colormap and the activity component with hot colormap are overlaid. The boundary of the region of interest (ROI) is drawn with a white dotted line. Five single pixels to be analyzed are marked with cyan arrows. **b**, Histogram of single pixel signal-to-noise ratio (SNR) from raw data and SUPPORT denoised data. **c**, Traces extracted from SUPPORT denoised data for the ROI in **a**, electrophysiological recording, injected currents. **d**, Traces extracted from raw and SUPPORT denoised videos from single pixels in **a** and electrophysiological recording. Left: From raw video. Right: From SUPPORT denoised video.

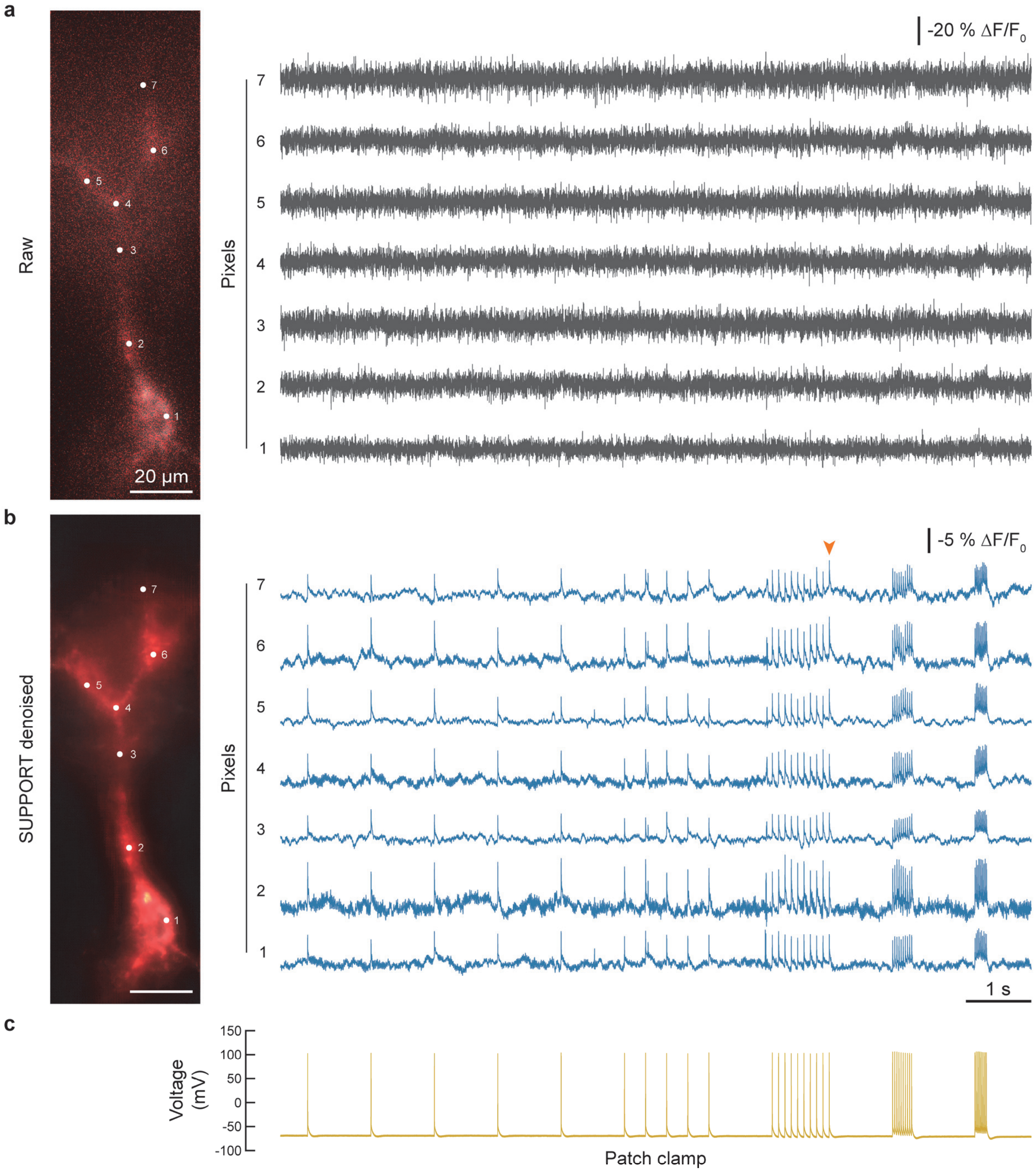


**Supplementary Fig. 4: SUPPORT reveals the voltage signal along the axonal branch. a**, Left: A representative frame of raw video after baseline correction. The frame used is marked on **b** with an orange arrow. Voltron2-expressing mouse cortex L2/3 was used as a dataset (Supplementary Table 1). The baseline component with gray colormap and the activity component with hot colormap are overlaid. Single pixels to be analyzed along the axonal branch are marked with white dots and numbers. Right: Traces extracted from the single pixels of raw video. **b**, Left: A representative frame of SUPPORT denoised video after baseline correction. Right: Traces extracted from the single pixels of SUPPORT denoised video. **c**, Corresponding voltage traces from the electrophysiological recording.

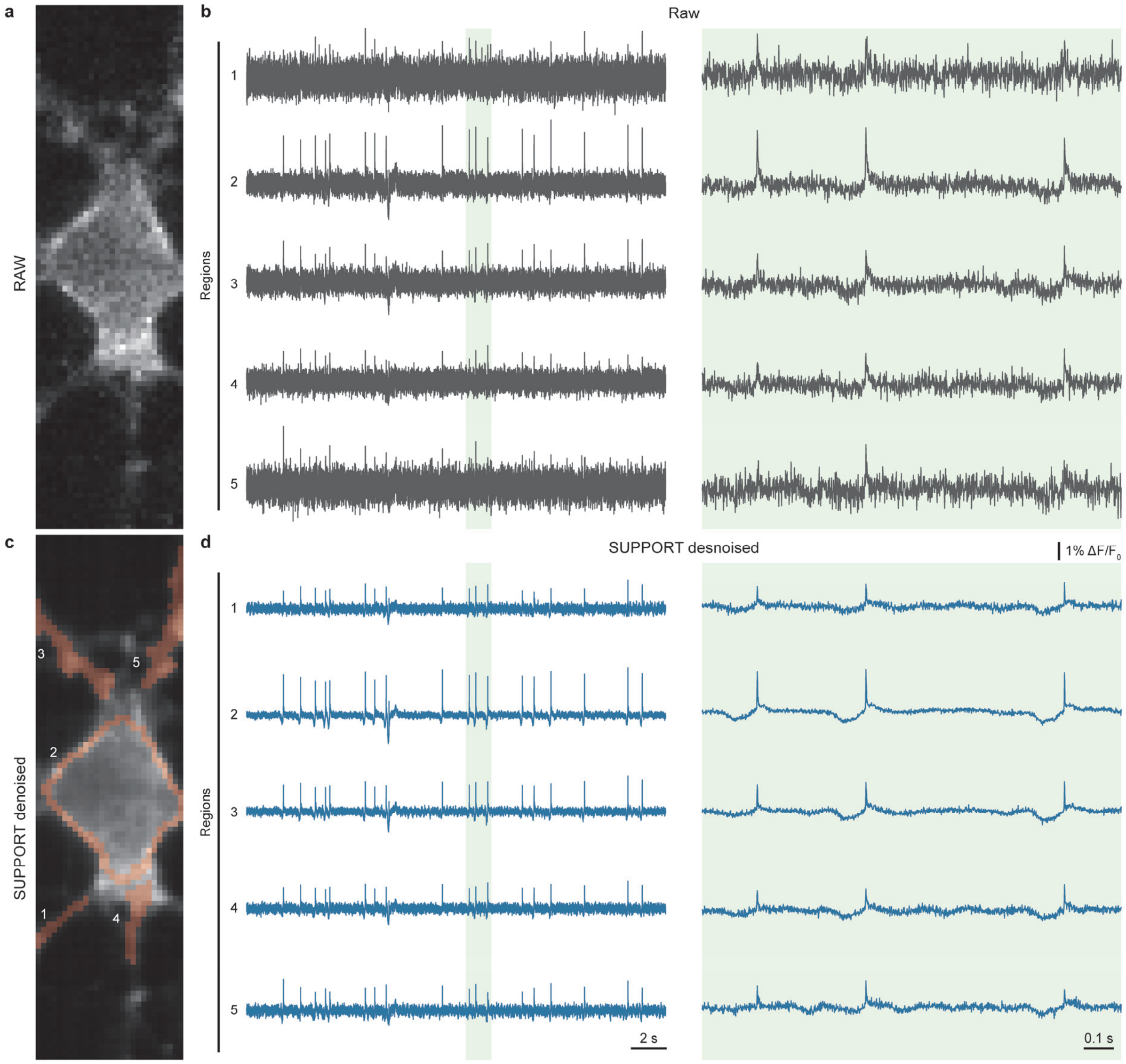


**Supplementary Fig. 5: SUPPORT reveals the voltage signal for several regions. a**, Representative frame of raw video. BeRST1-labeled mouse hippocampus cultured cells were used as a dataset (Supplementary Table 1). **b**, Traces extracted from the five regions of interest (ROIs) shown in **c** from the raw video. Temporally expanded traces from the green area on the left are shown on the right. **c**, Representative frame of SUPPORT denoised video. ROIs to be analyzed are marked in orange. **d**, Left: Traces extracted from the five ROIs shown in **c** from the SUPPORT denoised video. Right: Temporally expanded traces from the green area on the left.

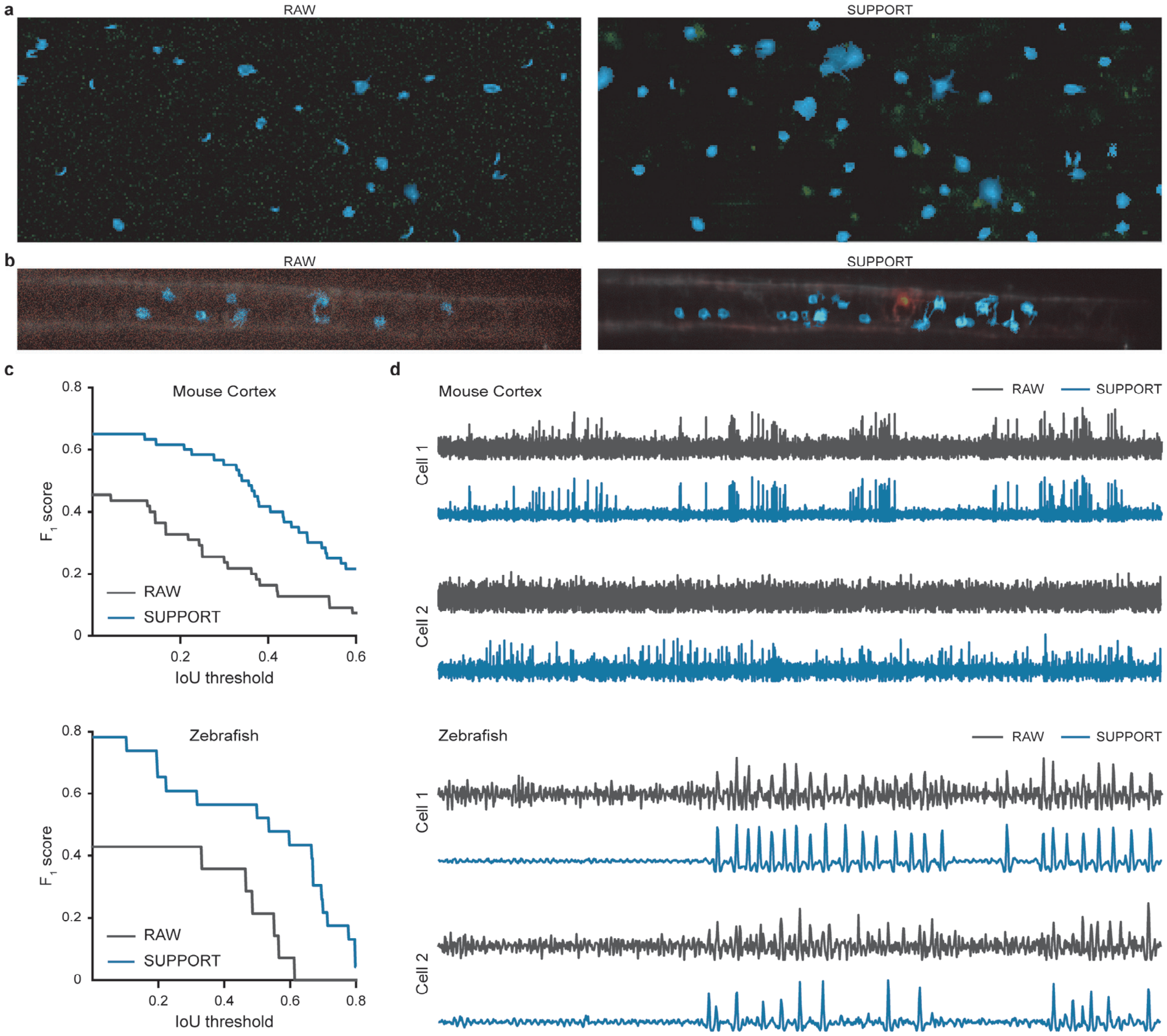


**Supplementary Fig. 6: Neuronal source extraction using localNMF from population voltage imaging. a**, Extracted neurons from raw and SUPPORT denoised mouse videos are colored in blue and overlaid on the images of Fig. 4a (Methods). **b**, Extracted neurons from raw and SUPPORT denoised zebrafish videos are colored in blue and overlaid on the images of Fig. 4d. (Methods) **c**, $F_1$ scores across intersection-over-union thresholds. **d**, Extracted temporal signals from raw and SUPPORT denoised video for two representative cells.

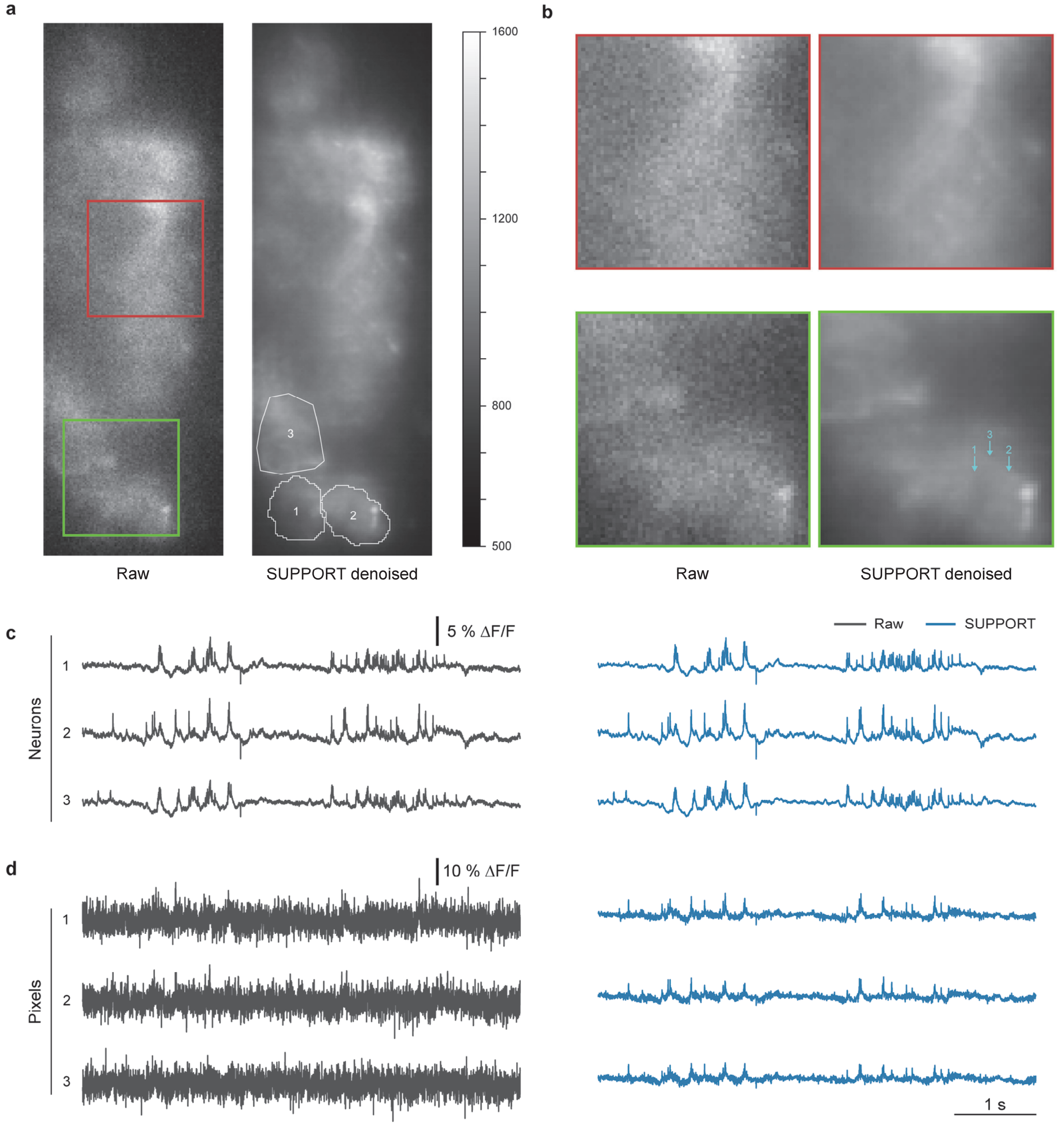


**Supplementary Fig. 7: Applying SUPPORT to population voltage imaging data with paQuasAr3s indicator. a**, Representative frames from raw video and SUPPORT denoised video. paQuasAr3s-expressing mouse hippocampus CA1 was used as a dataset (Supplementary Table 1). Boundaries of three regions of interest (ROIs) are drawn with white lines. **b**, Magnified views of the boxed regions in **b**. **c**, Traces extracted from raw and SUPPORT denoised videos for three ROIs in **a**. **d**, Single pixel traces from raw and SUPPORT denoised video for three pixels indicated in **b**.

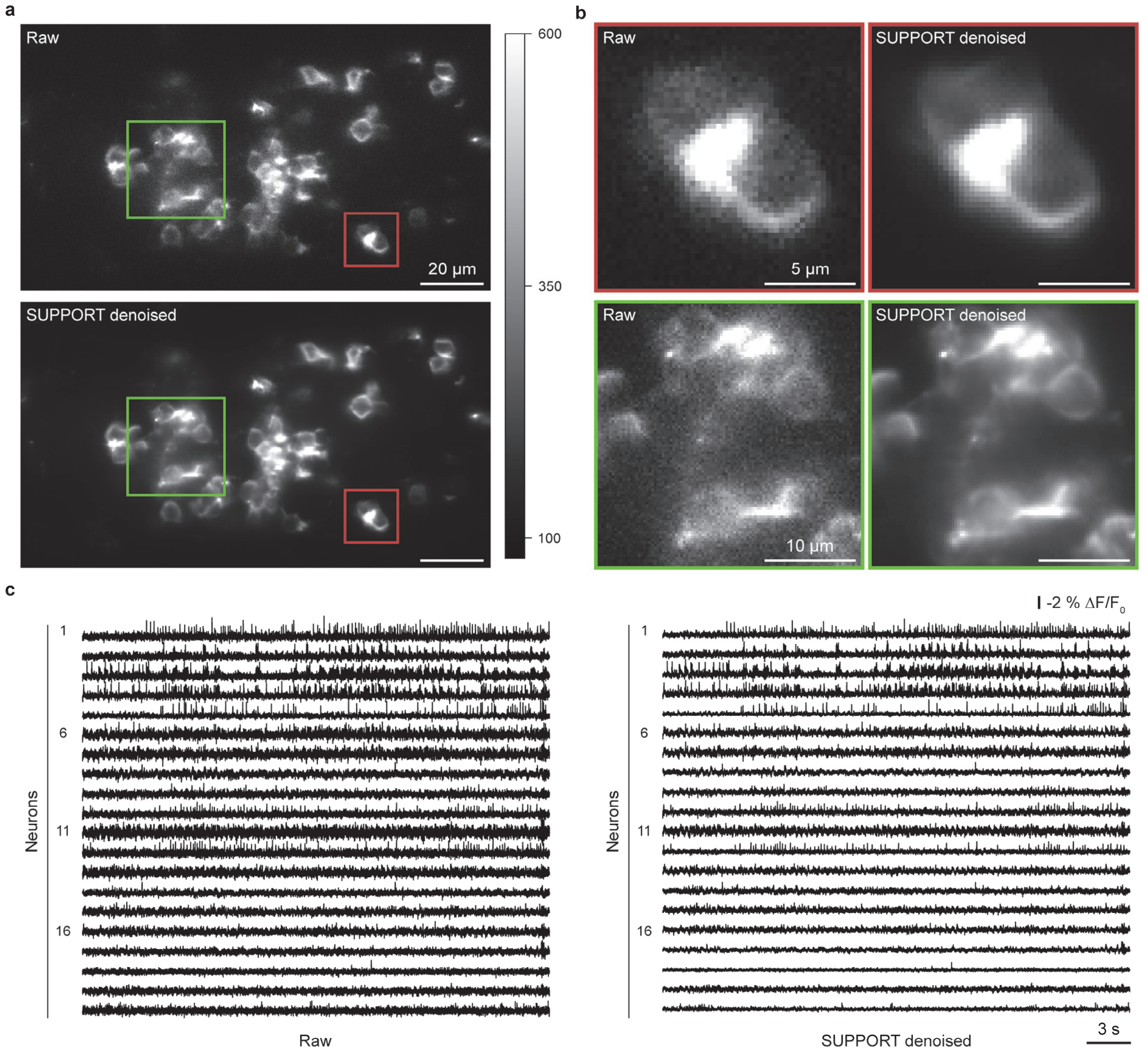


**Supplementary Fig. 8: Applying SUPPORT to population voltage imaging data with Voltron1 indicator. a**, Representative frames from raw video and SUPPORT denoised video. Voltron1-expressing zebrafish tegmental area was used as a dataset (Supplementary Table 1). **b**, Magnified views of the boxed regions in **b**. **c**, Traces extracted from the raw and SUPPORT denoised video for 20 neurons. Left: From the raw video. Right: From SUPPORT denoised video.

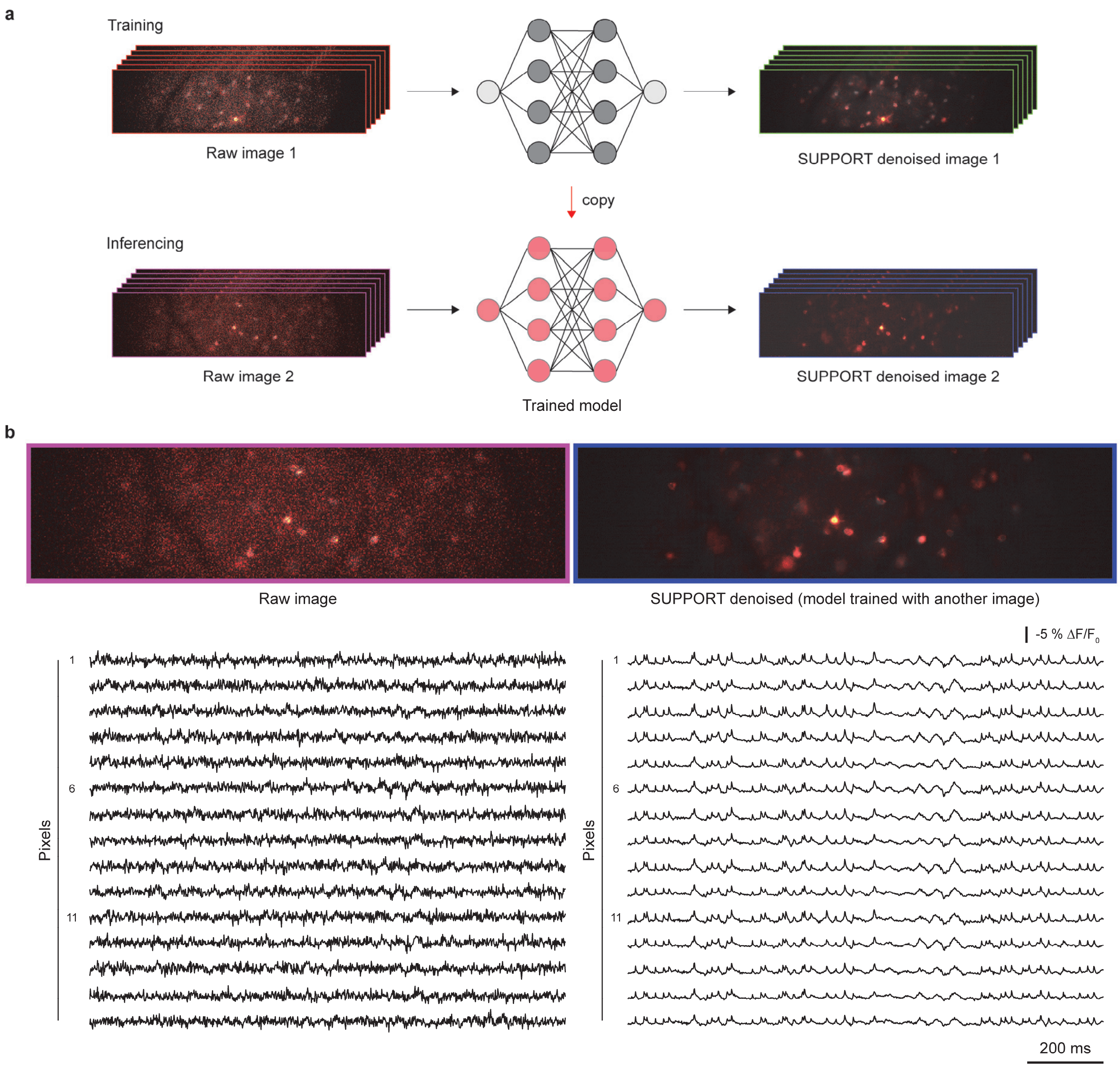


**Supplementary Fig. 9: Network generalization for population voltage imaging data. a**, Ability to generalize SUPPORT for denoising unseen dataset. Top: SUPPORT is first trained with raw image 1. Bottom: Trained SUPPORT network is used for denoising raw image 2, which is recorded from different fields of view and time. **b**, SUPPORT was able to denoise the unseen dataset without additional fine-tuning. Top: A representative frame of raw video and SUPPORT denoised video, for which the raw data was never shown to SUPPORT on training. Bottom left: Single pixel traces inside one neuron from the raw video. Bottom right: Traces from SUPPORT denoised video. This shows that the traces from SUPPORT denoised video are highly correlated.

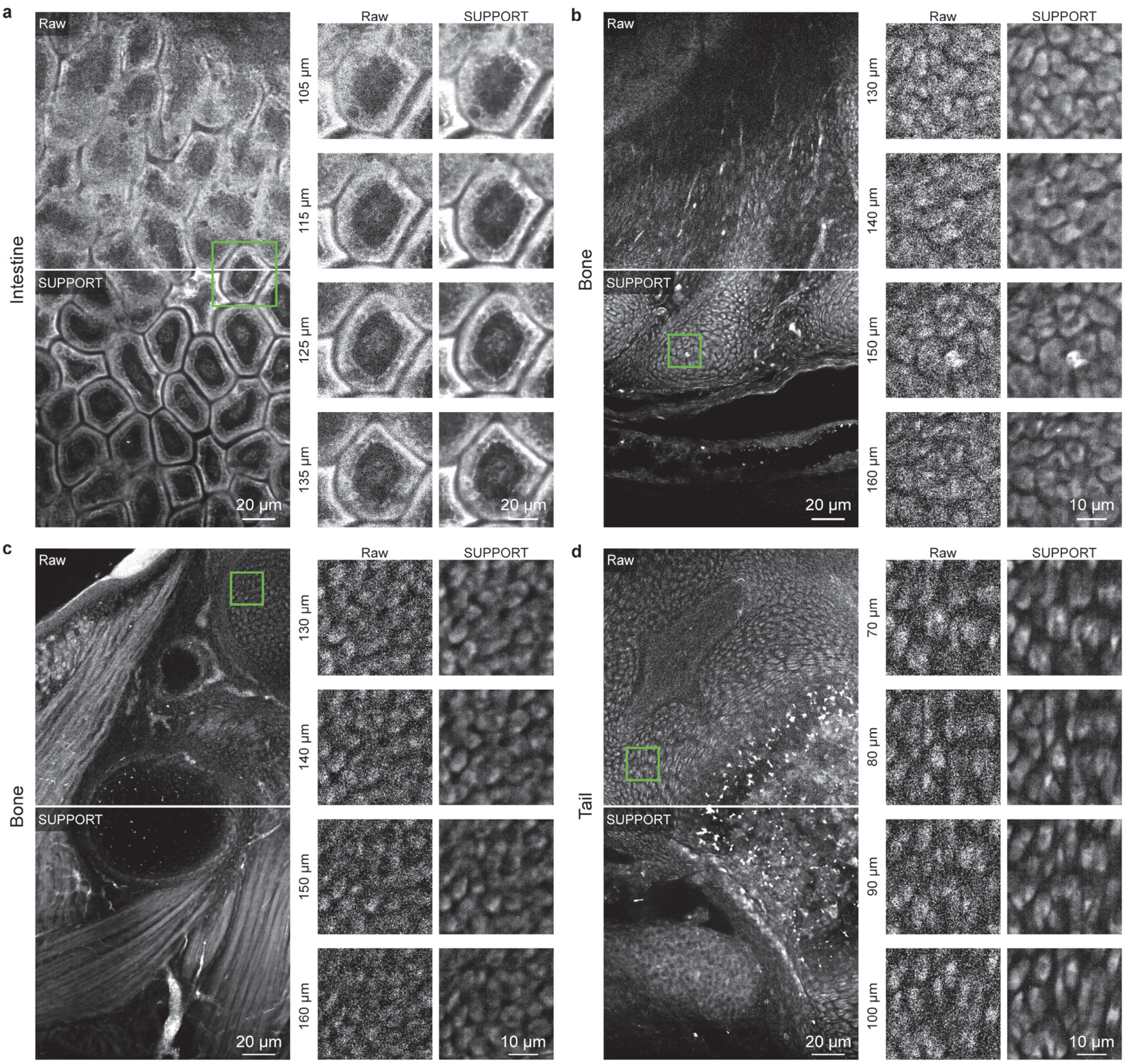


**Supplementary Fig. 10: Applying SUPPORT to volumetric structural images of mouse embryos acquired with expansion microscopy. a–d**, Left: A frame at depth 125 µm from raw data (top) and SUPPORT denoised data (bottom). Right: Expanded view of green box on the left at multiple depths. **a,** Intestine. **b-c**, Bone. **d**, Tail.

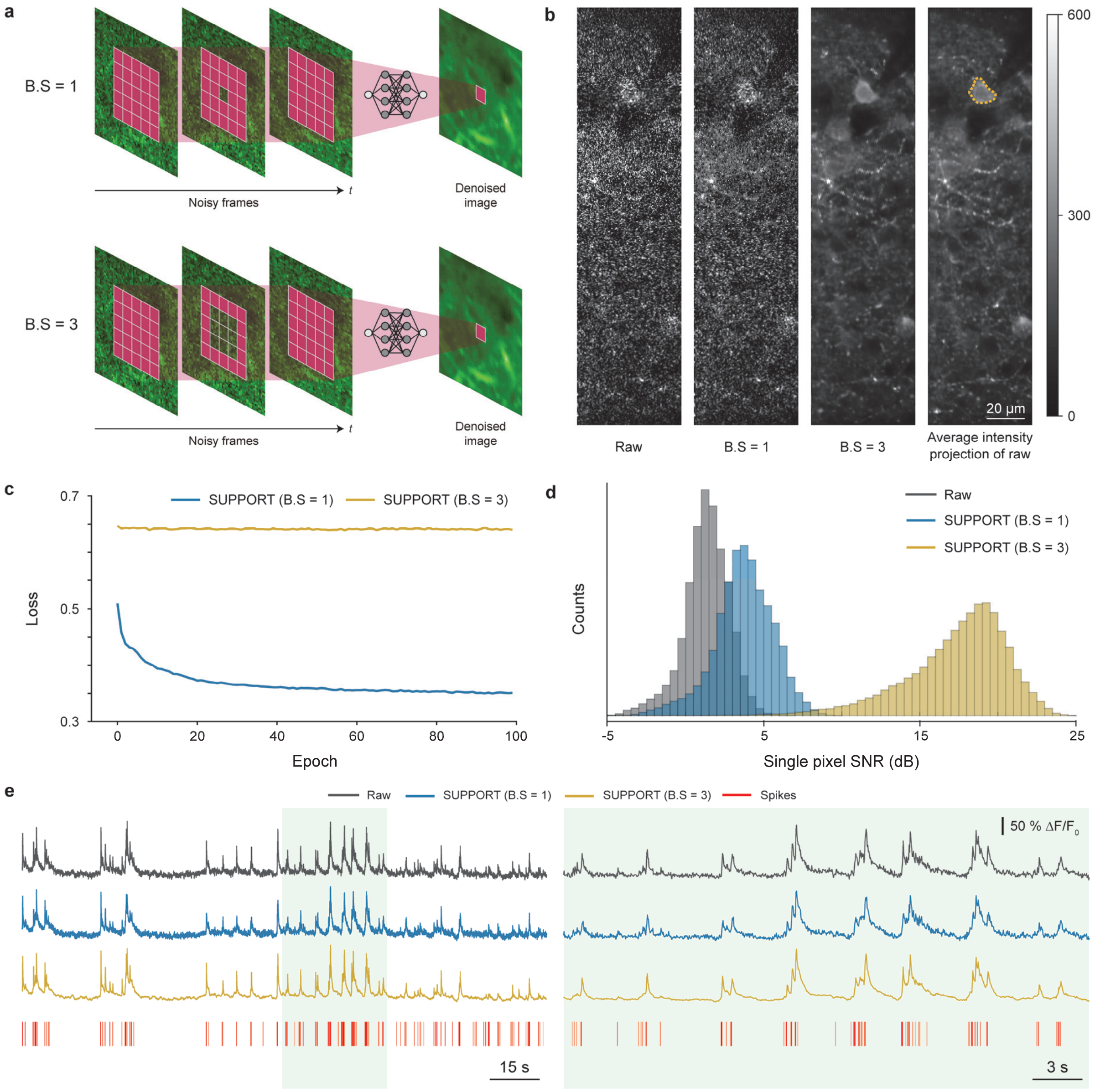


**Supplementary Fig. 11: Increasing the size of the blind spot to denoise the data with structured noise.** **a**, Receptive fields with different sizes of the blind spot. Top: Blind spot size of 1. Bottom: Blind spot size of 3. The center frame of noisy frames is the current frame to be denoised. **b**, Representative frames of raw and SUPPORT denoised data with structural noise. Raw data were recorded from a jGCaMP8f-expressing mouse cortex L2/3 (Supplementary Table 1). From left to right: Raw data, SUPPORT denoised data with blind spot sizes of 1 and 3, and the average intensity projection of the raw data. The boundary of the region of interest (ROI) is drawn with a yellow dotted line. **c**, Training loss curve of SUPPORT with blind spot sizes of 1 and 3. With a blind spot size of 1, SUPPORT learned to predict the structured noise, and training loss was significantly lower than for a blind spot size of 3. **d**, Histogram of single pixel signal-to-noise ratio (SNR) from raw data, SUPPORT denoised data with blind spot sizes of 1 and 3. While a blind spot size of

3 significantly increased the SNR, a size of 1 only slightly increased the SNR. **e**, Traces extracted from the raw data, SUPPORT denoised data with blind spot sizes of 1 and 3 for the ROI in **b**, and spikes (action potentials) detected from simultaneous electrophysiological recording. Temporally expanded traces from the green area of the left are shown on the right.

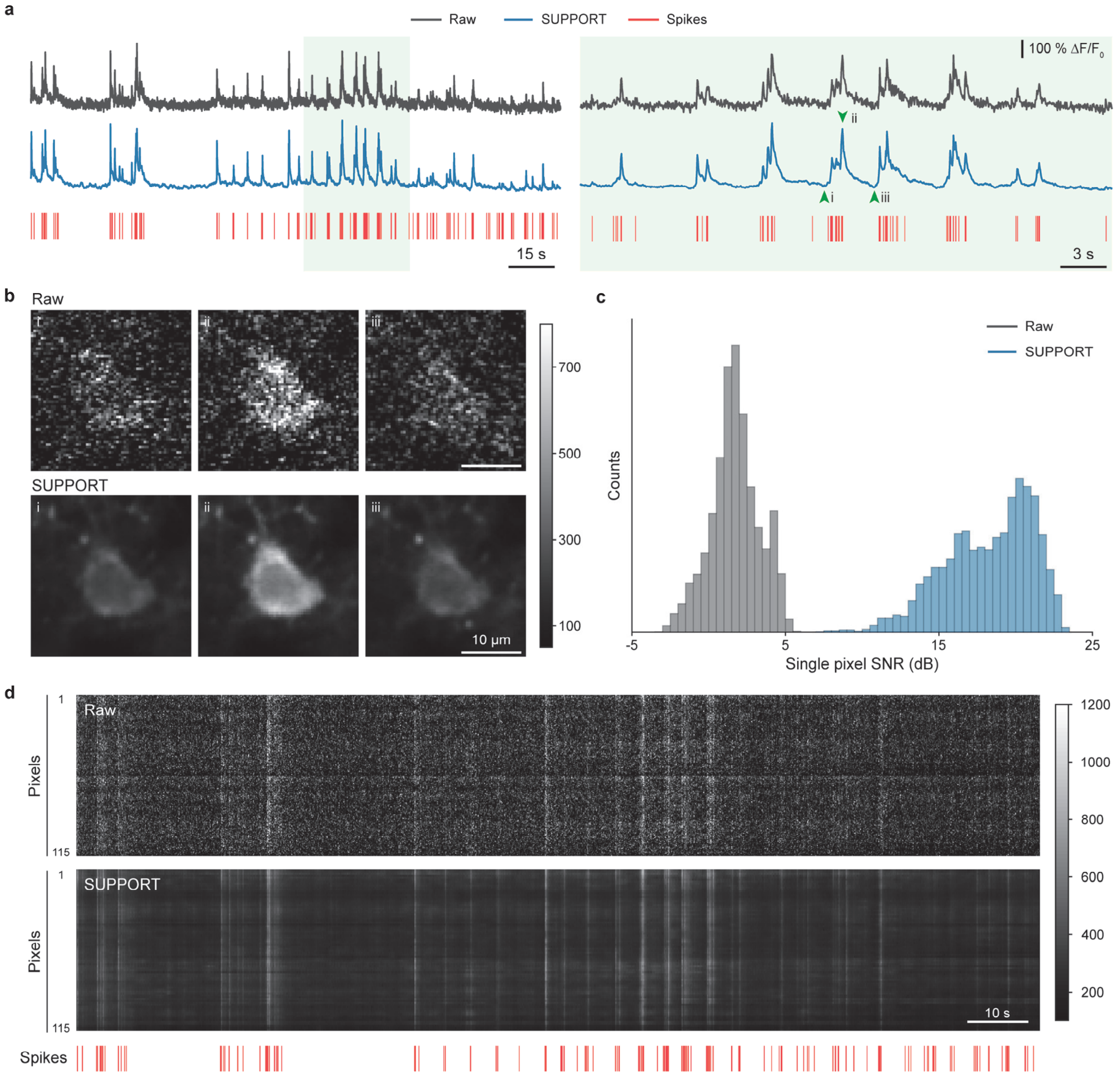


**Supplementary Fig. 12: Applying SUPPORT to simultaneous calcium imaging and electrophysiological recording data with jGCaMP8f indicator. a**, Traces extracted from the cytoplasmic area of raw and SUPPORT denoised video. Raw data were recorded from a jGCaMP8f-expressing mouse cortex L2/3 (Supplementary Table 1). Spikes (action potentials) detected from simultaneous electrophysiological recordings are drawn at the bottom. Temporally expanded traces from the green area of the left are shown on the right. **b**, Representative frames of the raw and SUPPORT denoised video marked with green arrows on **a**. Top: From the raw video. Bottom: From the SUPPORT denoised video. **c**, Histogram of single pixel signal-to-noise ratio (SNR) from the raw video and SUPPORT denoised video. **d**, Traces for pixels in the cell from raw and denoised data. Each row corresponds to a signal of each pixel. Spikes detected from simultaneous electrophysiological recordings are plotted underneath.

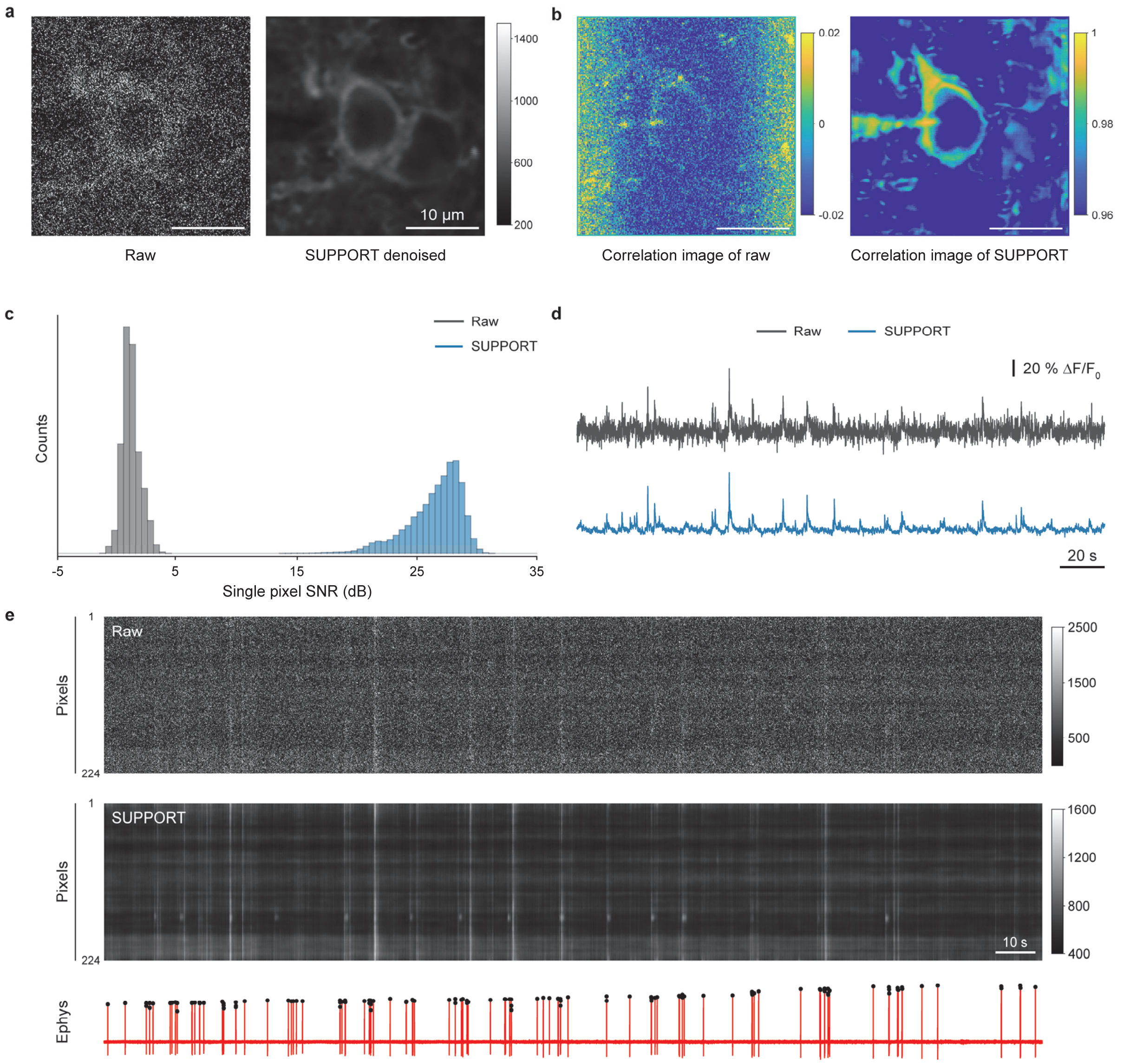


**Supplementary Fig. 13: Applying SUPPORT to simultaneous calcium imaging and electrophysiological recording data with GCaMP6f indicator. a**, Representative frames of raw video and SUPPORT denoised video. Raw data were recorded from a GCaMP6f-expressing mouse cortex V1 (Supplementary Table 1). **b**, Correlation images of raw video and SUPPORT denoised video. The correlation image for each pixel is the average of the correlation coefficients between the signal of that pixel and the signals of neighboring pixels. **c**, Histogram of single pixel signal-to-noise ratio (SNR) from raw and SUPPORT denoised videos. **d**, Traces extracted from the ROI of raw and SUPPORT denoised videos. The ROI contains 100 pixels inside the cytoplasm. **e**, Traces for pixels in the cell from raw and SUPPORT denoised video. Each row corresponds to a signal of each pixel. Simultaneous electrophysiological recording is plotted underneath, with black dots for detected spikes.

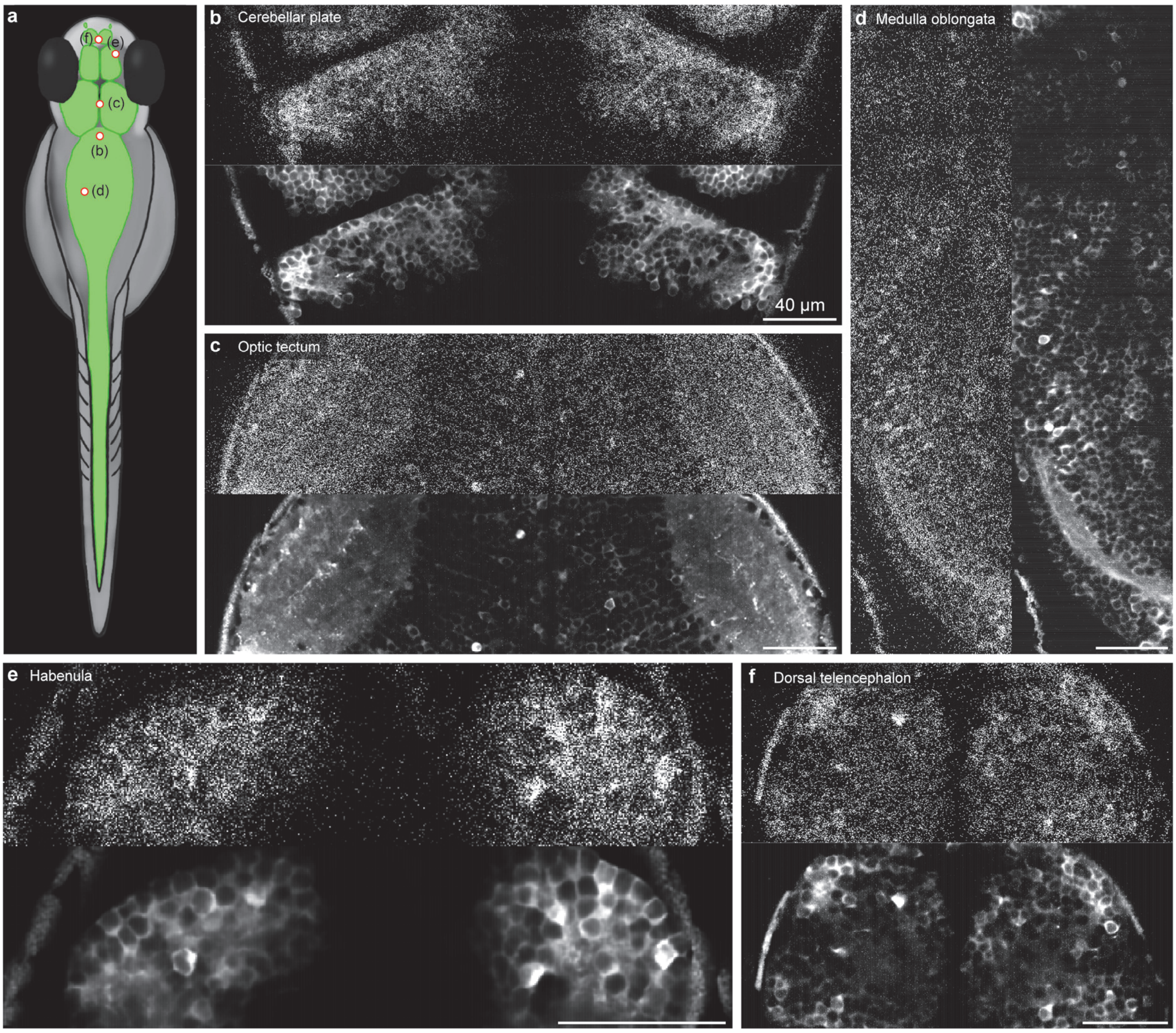


**Supplementary Fig. 14: SUPPORT denoises calcium imaging in larval zebrafish.** **a**, Larval zebrafish expressing GCaMP7a calcium indicator under control of *huc* promoter. **b–f**, Top left: Representative frames of raw video. Bottom right: Representative frame of SUPPORT denoised video. **b**, Cerebellar plate. **c**, Optic tectum. **d**, Medulla oblongata. **e**, Habenula. **f**, Dorsal telencephalon.

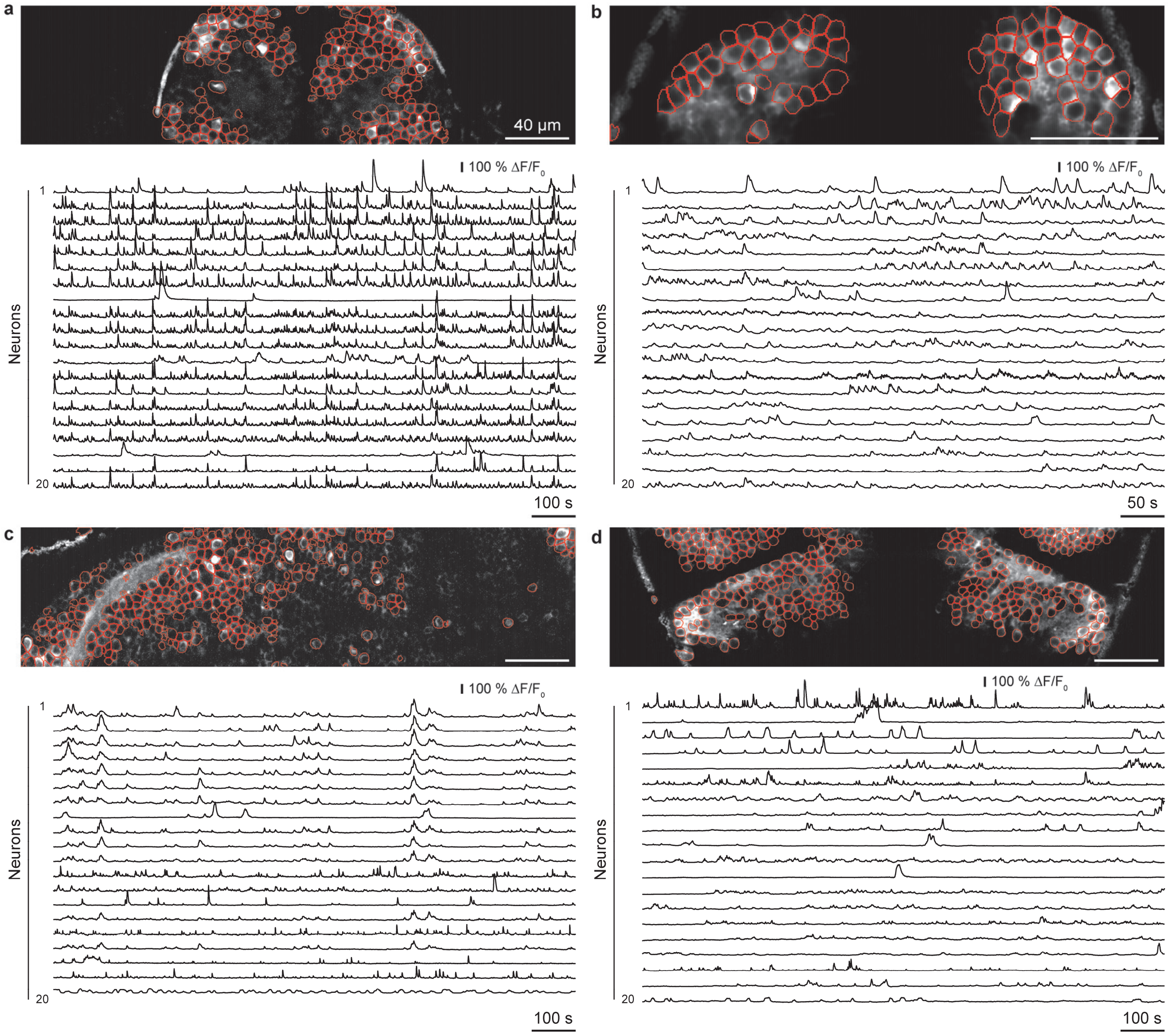


**Supplementary Fig. 15: SUPPORT denoising enables single frame cell detection. a–d**, Top: Neurons extracted from a single frame of SUPPORT denoised video. Bottom: Representative calcium traces of 20 neurons. **a**, Dorsal telencephalon. **b**, Habenula. **c**, Medulla oblongata. **d**, Cerebellar plate.

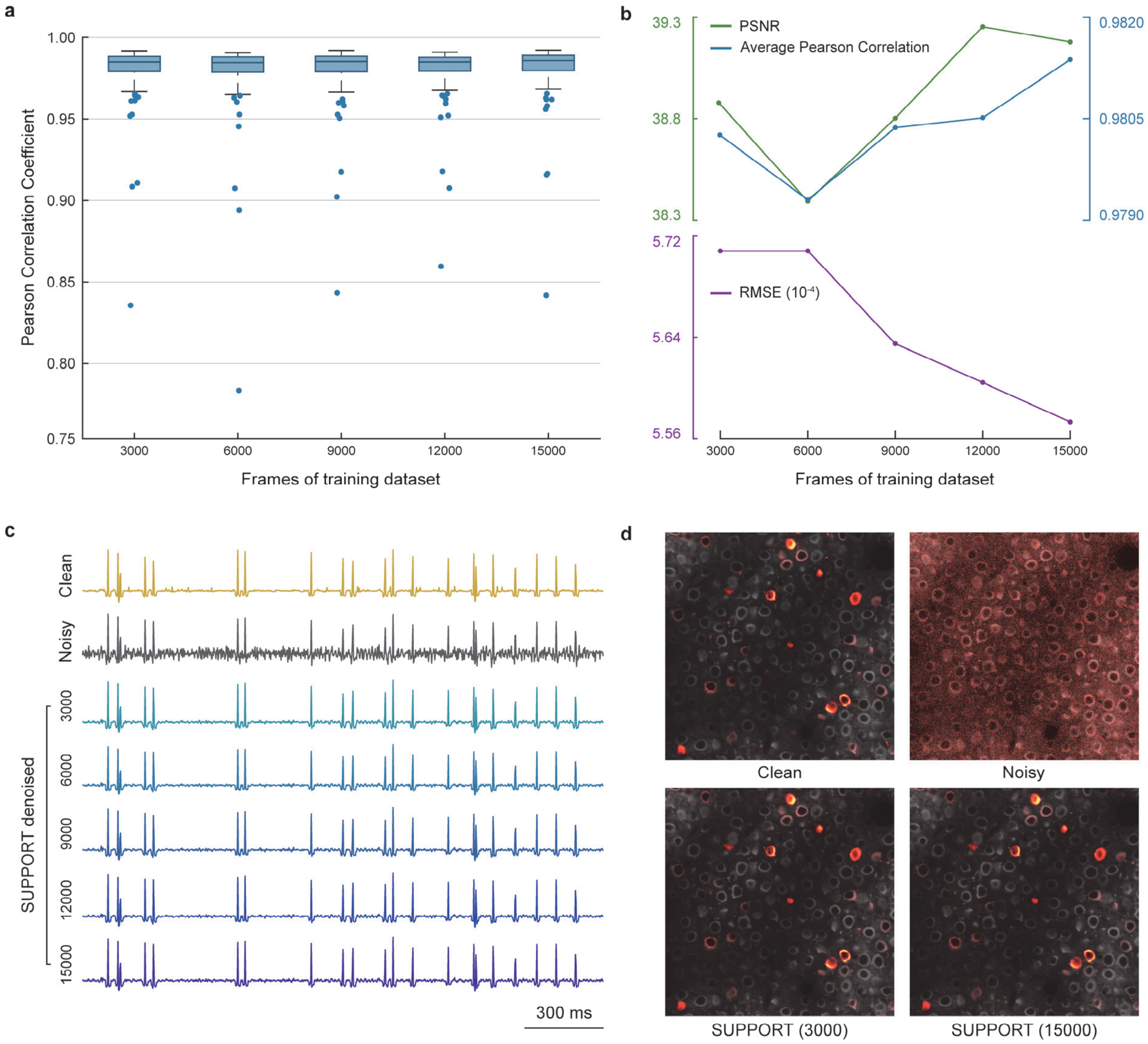


**Supplementary Fig. 16: Denoising performance as a function of the size of training data. a**, Pearson correlation coefficients between trace extracted from clean (ground truth) video and SUPPORT denoised videos. Synthetic data with spike width of 3 ms were used for comparison. SUPPORT was trained with five different sizes of training datasets to evaluate the correlation between denoising performance and dataset size. **b**, PSNR, average Pearson correlation coefficients, and RMSE between clean video and SUPPORT denoised videos. All three metrics show a trend of performance increases with larger training datasets. **c**, Traces extracted from a single cell of the clean video, SUPPORT denoised videos, and the noisy video. While performance improves with larger datasets, traces show that SUPPORT was able to denoise with 3,000 frames of training data. **d**, Representative frames of clean video, noisy video, and SUPPORT denoised videos after baseline correction (Methods). SUPPORT trained with 3,000 frames and 15,000 frames were used. Representative frames of clean and SUPPORT denoised videos.

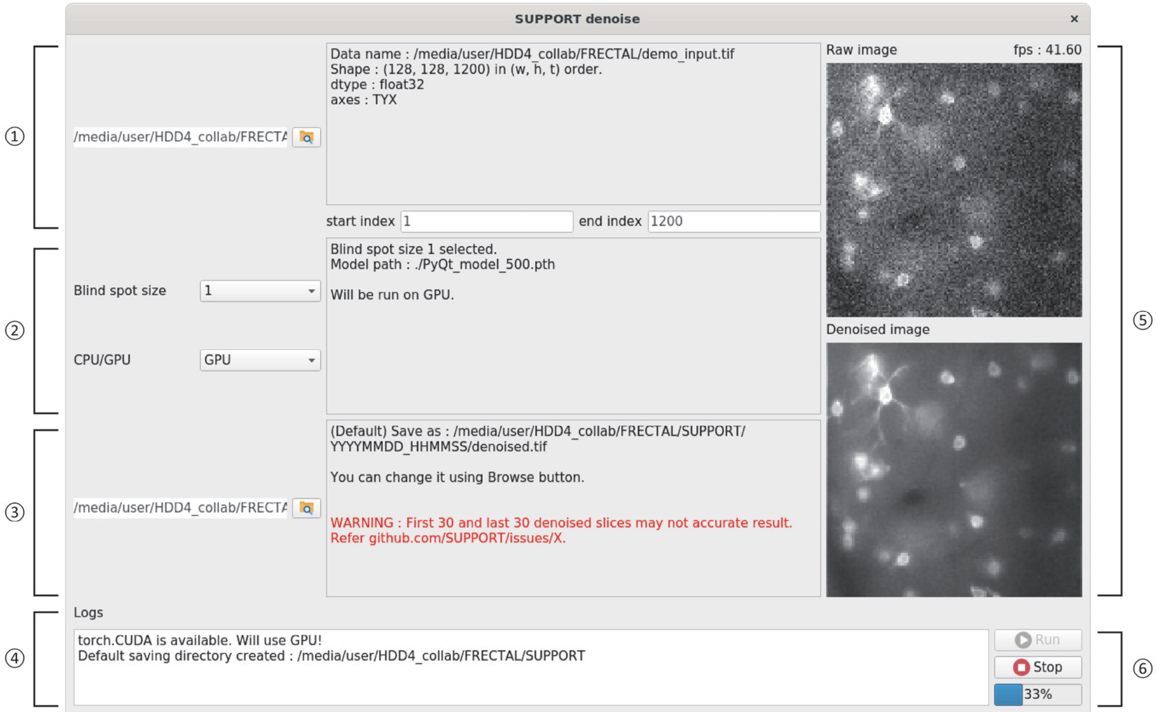


① (Input) Image path and information
② (Model) Blind spot size and CPU/GPU setting
③ (Save) Saving directory for denoised image and log files

④ Logs
⑤ Raw and denoised image in real time
⑥ Run / Stop button and progress bar

**Supplementary Fig. 17: Python GUI: SUPPORT software with a graphical user interface.** The user can choose the size of the blind spot and CPU/GPU option. Once the Run button is pressed, the software loads and denoises using a selected model, shows the denoised frame in real time, and saves denoised frames.

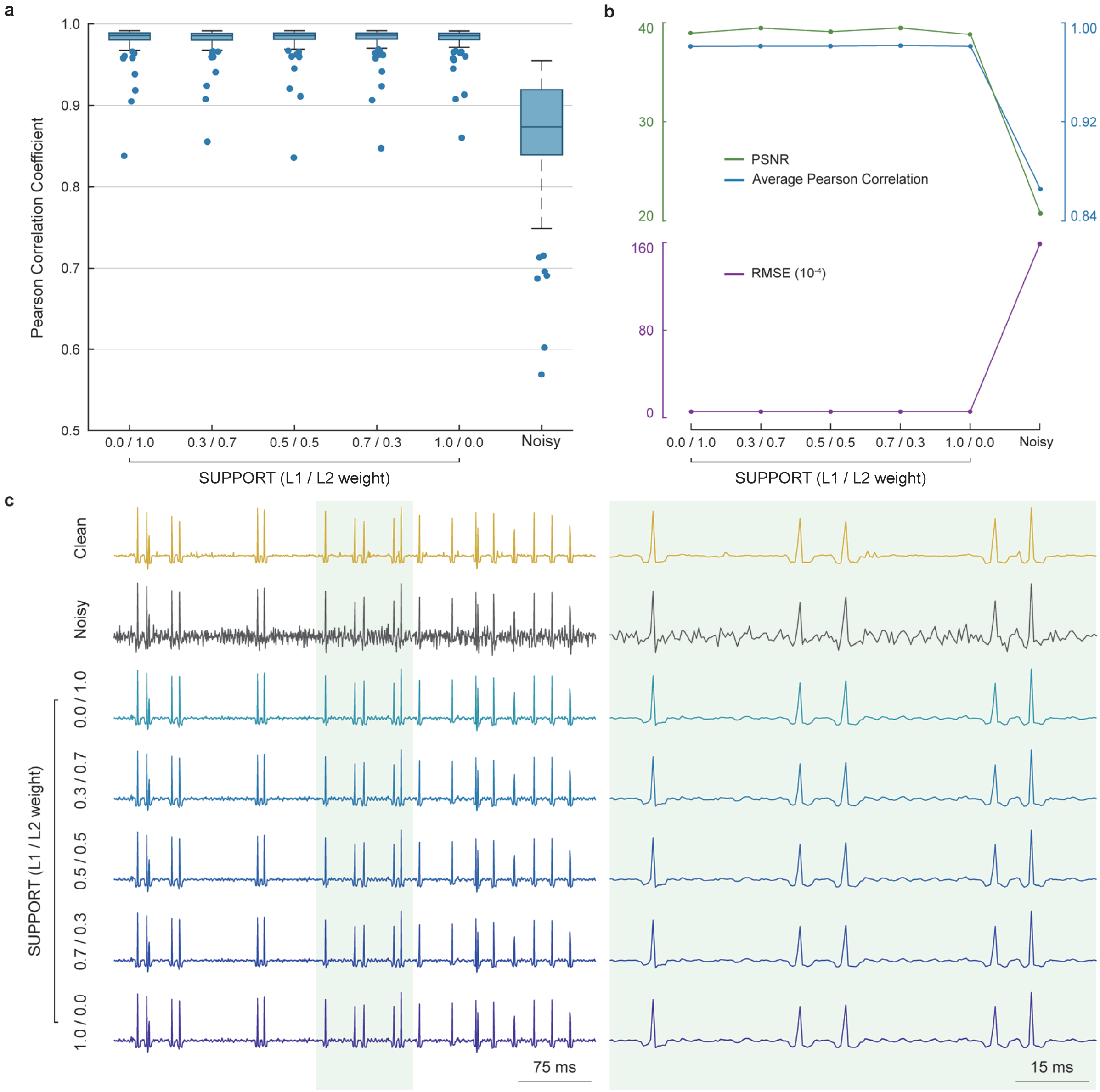


**Supplementary Fig. 18: L1 and L2 loss comparison with simulation data. a**, Pearson correlation coefficients between traces from clean (ground truth) video and SUPPORT denoised video or noisy video. SUPPORT denoised videos were acquired with five different training loss settings. **b**, PSNR, average Pearson correlation coefficients, and RMSE between clean video and SUPPORT denoised videos or noisy video. All three metrics indicate that the weight of L1 and L2 loss for training SUPPORT did not significantly change denoising performance. **c**, Traces extracted from a single cell of the clean video, SUPPORT denoised videos, and noisy video. Temporally expanded traces from the green area of the left are shown on the right. This again shows that the weight of L1 and L2 loss does not significantly change SUPPORT's output.

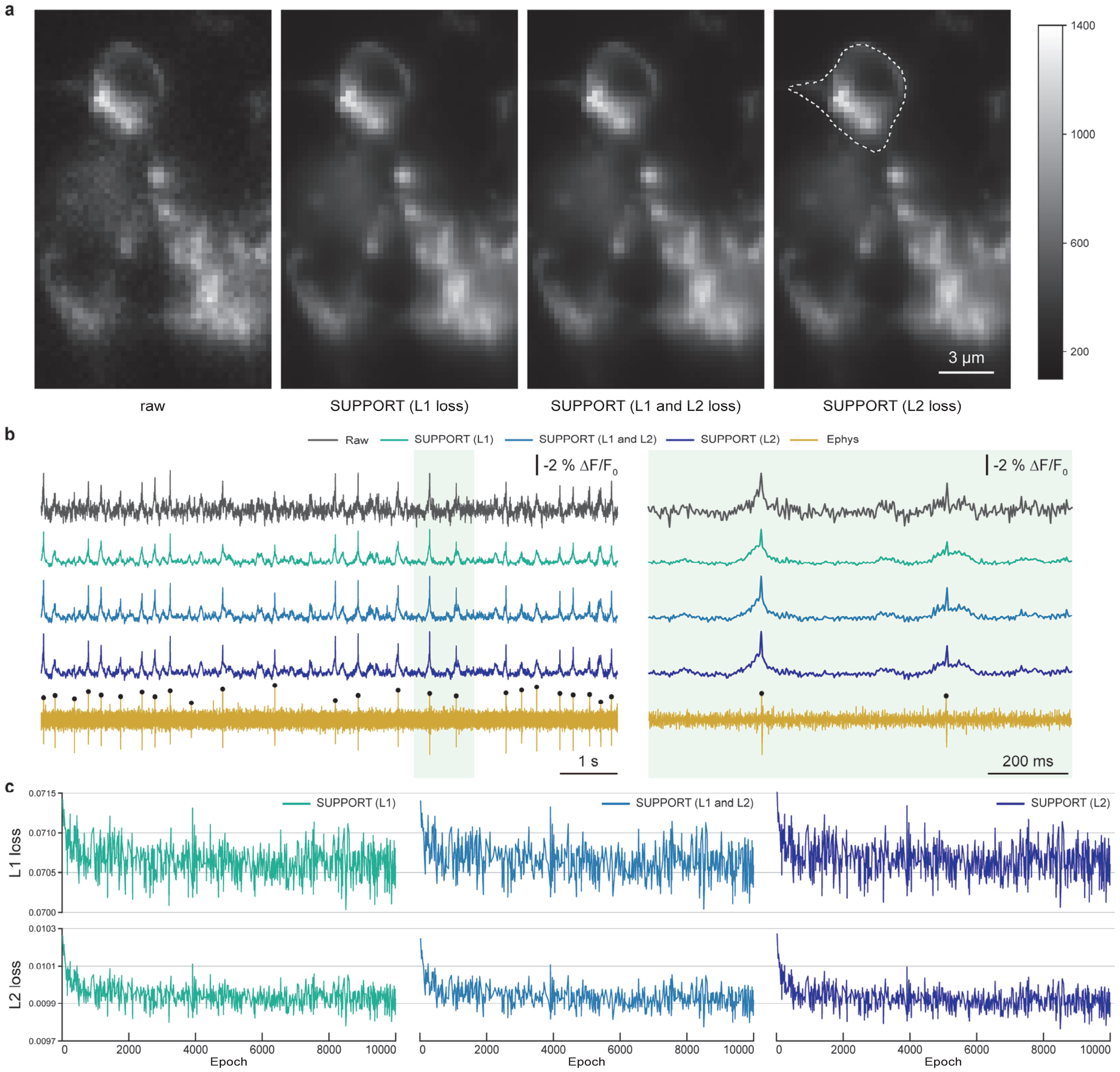


**Supplementary Fig. 19: L1 and L2 loss comparison with voltage imaging with simultaneous electrophysiological recording. a**, From left to right: Representative frames from the raw data, SUPPORT trained using L1 loss function, both L1 and L2 loss functions, and L2 loss function. For the training setting that used both L1 and L2 loss, equal weights of L1 and L2 were used. The boundary of the region of interest (ROI) is drawn with a white dotted line. **b**, Traces extracted from the raw data, SUPPORT trained using L1 loss function, both L1 and L2 loss function, and L2 loss function for the ROI in **a**, and electrophysiological recording. Detected spikes are marked with black dots. Temporally expanded traces from the green area of the left are shown on the right. Using additional L2 loss showed better reconstruction of the spikes after SUPPORT denoising. **c**, L1 loss and L2 loss were tracked during the training procedure for SUPPORT using L1 loss function, both L1 and L2 loss functions, and L2 loss function.

#### SUPPLEMENTARY VIDEOS

**Supplementary Video 1**. **Synthetic voltage imaging**

**Supplementary Video 2**. **Single-neuron simultaneous electrophysiology and voltage imaging of larval zebrafish**

**Supplementary Video 3**. **Population voltage imaging of larval zebrafish spinal cord**

**Supplementary Video 4**. **Volumetric time-lapse imaging of *C. elegans***

**Supplementary Video 5. Volumetric structural imaging of mouse embryos recorded with expansion microscopy**

**Supplementary Video 6**. **Calcium imaging of larval zebrafish**

#### SUPPLEMENTARY TABLES

| Region | Cell type | Recording rate (Hz) | Reporter | Imaging modality | Reference |
|---|---|---|---|---|---|
| **Voltage Imaging** | | | | | |
| Mouse cortex L1 (Fig. 1, 4, Supplementary Fig. 6, 9) | interneurons (*Ndnf*+) | 400 | Voltron1 | wide-field fluorescence | reporter and data: Abdelfattah, A. S. et al. [2] |
| Zebrafish dorsal part of the cerebellum (Fig. 3, Supplementary Fig. 19, Supplementary Video 2) | excitatory (*vGlut2a*) | 300 | Voltron1 + cell-attached extracellular recording | light sheet | reporter and data: Abdelfattah, A. S. et al. [2] |
| Zebrafish spinal cord (Fig. 4, Supplementary Fig. 6, Supplementary Video 3) | excitatory (*vGlut2a*) | 1000 | zArchon1 | light sheet | reporter: Piatkevich, K. D. et al. [3]<br>data: Xie, M. E. et al.[4] |
| Mouse cortex L2/3 (Supplementary Fig. 3) | pyramidal cells | 1000 | QuasAr6a + patch clamp | one-photon epifluorescence microscopy with targeted illumination | reporter: Tian, H. et al.[5]<br>data: this work |
| Mouse cortex L2/3 (Supplementary Fig. 4) | pyramidal cells | 1000 | Voltron2 + patch clamp | one-photon epifluorescence microscopy with targeted illumination | reporter: Abdelfattah, A. S. et al.[6]<br>data: this work |
| Cultured rat hippocampal neuron (Supplementary Fig. 5) | primary neurons | 1000 | BeRST1 | wide-field fluorescence | reporter: Huang, Y. L., Walker, A. S. & Miller, E. W.[7]<br>data: this work |

| | | | | | |
|---|---|---|---|---|---|
| Mouse hippocampus CA1 (Supplementary Fig. 7) | interneurons | 1000 | paQuasAr3-s | micromirror-based, soma-targeted, structured illumination | reporter and data: Adam, Y. et al.[8] |
| Zebrafish tegmental area (Supplementary Fig. 8) | excitatory (*vGlut2a*) | 300 | Voltron1 | light sheet | reporter and data: Abdelfattah, A. S. et al.[2] |
| **Structural Imaging** | | | | | |
| *C. elegans* (Fig. 5, Supplementary Video 4) | pan-neuronal (*H20*) | 4.75 | mCherry | confocal | reporter: Shaner, N. C. et al.[9] data: Toyoshima, Y. et al.[10] |
| *Penicillium* (Fig. 6) | entire | Static | N/A | confocal | data: this work |
| Mouse embryos; intestine, bone, tail (Fig. 6, Supplementary Fig. 10) | entire | Static | Alexa Flour 488 NHS-ester | confocal | data: Sim, J. et al.[11] |
| **Calcium Imaging** | | | | | |
| Mouse cortex L2/3 (Supplementary Fig. 11, 12) | pyramidal cells | 122 | jGCaMP8f + loose-seal cell attached electrophysiological recording | two-photon | reporter: Zhang, Y. et al.[12] data: Rozsa, M. et al.[13] |

| | | | | | |
|---|---|---|---|---|---|
| Mouse cortex V1 (Supplementary Fig. 13) | excitatory (*syn*) | 60 | GCaMP6f + loose-seal cell attached electrophysiological recording | two-photon | reporter: Chen, T. W. et al.[14]<br>data: GENIE project[15] |
| Zebrafish (Supplementary Fig. 14, 15, Supplementary Video 5) | pan-neuronal (*huc*) | 1, 2, 4 | GCaMP7a | confocal | reporter: Muto, A., Ohkura, M., Abe, G., Nakai, J. & Kawakami, K.[16]<br>data: this work |

**Supplementary Table 1. List of datasets analyzed in this study.**
